# Vestibulomotor Weighting Associated with Cybersickness in Virtual Reality

**DOI:** 10.64898/2026.05.04.722436

**Authors:** Megan H. Goar, Michael Barnett-Cowan

## Abstract

Cybersickness is a major barrier to the widespread adoption of virtual reality (VR), yet its underlying neurophysiological mechanisms remain poorly understood. This study investigated the relationship between vestibulomotor weighting and cybersickness. Vestibulomotor weighting was quantified using electrical vestibular stimulation (EVS), with coherence and gain between the EVS input and medial-lateral center-of-pressure (ML-CoP) responses indexing the contribution of vestibular input to postural control. Thirty-eight healthy young adults (females n=21, males n=17) completed a standing VR rollercoaster task while receiving continuous stochastic EVS (0–25 Hz; ±4.5 mA), with ML-CoP responses recorded using a force plate.

Cybersickness was assessed using the Fast Motion Sickness Scale (FMS) and Simulator Sickness Questionnaire, and participants were classified as non-sick (FMS < 5), medium-sick (FMS ≥ 5), or high-sick (terminated the VR exposure early due to intolerance). Baseline EVS-ML-CoP coherence across 2.5–8 Hz was significantly greater in high-sick than in non-sick participants, indicating elevated vestibulomotor weighting in individuals who developed symptoms. During VR exposure, coherence declined over time in symptomatic groups (mean slope = −0.0027 for medium-sick), whereas non-sick participants maintained consistently low coherence (mean slope = −0.0005). Despite this reduction in vestibular coupling, postural sway increased in the high-sick group relative to the medium-and non-sick groups (+29% vs. −7% and −30% change in ML-CoP RMS, respectively), while vestibular-evoked response amplitude decreased (gain reduced by 64% across 2.5–3.5 Hz). These findings indicate that greater baseline vestibulomotor weighting was associated with increased susceptibility to cybersickness, whereas reductions in vestibular contributions during VR with EVS reflected adaptive reweighting that was insufficient to prevent instability and symptom progression. Together, the results highlight baseline sensory reliance as a key determinant of cybersickness vulnerability and suggest that reweighting during exposure plays a secondary, mitigating role.

**New and Noteworthy:** We provide the first evidence that baseline vestibulomotor weighting predicts susceptibility to cybersickness in virtual reality and is dynamically reduced during exposure. Using electrical vestibular stimulation, we show that symptomatic individuals begin with greater reliance on vestibular input for postural control and progressively downweight these signals in response to sensory conflict.

## Introduction

The central nervous system (CNS) maintains upright orientation and postural stability by integrating vestibular, visual, and somatosensory inputs (1). Sometimes, sensory conflict arises when mismatches occur between expected and actual afferent signals across sensory modalities (1–5). This sensory conflict is a primary driver of motion sickness (1,6), which is characterized by symptoms such as nausea, headaches, and vomiting (12–18). A visually induced form of motion sickness, called cybersickness, can occur when virtual environments simulate self-motion without corresponding vestibular or proprioceptive cues, and is often exacerbated by factors such as display latency (10,13,14). Despite the potential of virtual reality (VR) for applications in training, rehabilitation, and education, cybersickness remains a major barrier to its widespread use (10,11,13–15).

Susceptibility to cybersickness in VR varies substantially across individuals, with some tolerating prolonged VR exposure and others developing severe symptoms rapidly (16,17). It has been proposed that this may reflect differences in neurophysiological adaptation and sensory integration, particularly sensory reweighting (16,18–20). Evidence suggests that the magnitude of sensory reweighting is inversely proportional to cybersickness susceptibility (16,20).

However, it remains unclear which sensory cues should be prioritized during VR. Moreover, direct measurement of sensory weights and their dynamic adjustment remain technically challenging.

One method to measure sensory weights is by artificially perturbing individual sensory modalities, using methods such as using tendon vibration (21,22), visual scene motion (20,23–27), and vestibular stimulation (28–30). The resulting motor responses to these cues can be analyzed using linear systems approaches to quantify how the CNS utilizes sensory cues for sensorimotor control. In a linear systems framework, coherence quantifies the frequency-dependent coupling between sensory input and motor output, while gain represents the sensitivity of the sensorimotor response (31,32). Increases in coherence or gain can be interpreted as evidence that the CNS has upweighted the corresponding sensory cue, whereas decreases indicate downweighting (1,2,4,20,33).

Visually evoked balance responses have been quantified using linear systems analyses by relating trajectories of visual motion presented on a surround projection to postural sway.

Individuals who exhibit greater sway at higher visual oscillation amplitudes (i.e., increased gain, reflecting greater visual reliance) are more likely to experience increased cybersickness symptom severity, suggesting that lower visual weighting may be advantageous (20).

While visual probes characterize the response to scene motion, stochastic electrical vestibular stimulation (EVS) enables the isolated probing of vestibular-evoked balance responses non-invasively. EVS activates vestibular afferents in the inner ear, which signal head motion (i.e., translation or rotation) and tilt relative to gravity (34). When combined with linear systems analyses, stochastic EVS allows for quantification of coherence and gain in balance responses, providing an objective measure of vestibular contributions to sensorimotor control (2,33,35–40).

Since EVS simultaneously activates afferents with varying directional sensitivities, the resulting neural signal is not fully congruent with natural head motion (28,30). At higher stimulation intensities (greater than 1 mA; (29)), EVS can therefore produce disorienting vestibular cues that disrupt balance and posture. In VR environments, EVS has the potential to both increase sensory conflict, thereby exacerbating cybersickness, and to serve as a quantitative probe of vestibular modulation relative to sickness severity.

Previous work has demonstrated that vestibular-evoked response gain increases when visual self-motion acuity is reduced (and vice versa) via LED-based displays (41). However, it remains unclear whether this relationship generalizes to immersive VR environments, where visual motion cues are more complex, multisensory conflict is heightened, and sickness can develop rapidly.

Evidence linking sensory weights and reweighting to cybersickness remains minimal, with studies often showing small effect sizes and substantial inter-individual variability. No clear relationship has been established between weighting strategy and cybersickness susceptibility.

Furthermore, traditional studies measure sensory reweighting only before and after VR exposure, potentially overlooking transient adaptations that occur during sickness development. The current study addresses this limitation by linking real-time vestibulomotor reweighting, quantified through coherence and gain between EVS input and mediolateral center of pressure (ML-CoP) output, to individual sickness trajectories during VR gameplay. This approach provides a novel window into the temporal dynamics of neural adaptation during cybersickness.

The objective of this study was to examine whether vestibulomotor weights differ between highly susceptible individuals and non-sick individuals, and whether these weights change over time. We induced strong visual-vestibular conflict using a passive rollercoaster VR game while recording ML-CoP responses to continuous EVS to characterize vestibulomotor reweighting. Based on previous findings that greater visual weighting is associated with increased cybersickness (20), we hypothesized that individuals who rely more heavily on vestibular input to compensate for unreliable visual cues would be more resilient to VR-induced sensory conflict. Specifically, we predicted that greater vestibulomotor coupling (EVS-ML-CoP coherence) and larger response amplitudes (gain) would be associated with reduced sickness severity. We further predicted that EVS-ML-CoP coherence and gain would change over time during VR exposure, reflecting sensory reweighting.

Identifying weighting profiles that predict cybersickness resilience will advance fundamental neuroscience by improving our understanding of multisensory adaptation. These insights may also guide targeted interventions to promote safe and inclusive use of VR in education, healthcare, and industry, especially for individuals currently excluded due to susceptibility.

## Methods

### Participants and Ethical Approval

A sample of 38 participants (females n=21, males n=17) was recruited. Sample size was determined a priori based on power calculations (n = 10 per group), with additional recruitment to account for variability in sickness responses. Young adults (18–35 yr old) with no self-reported neurological or orthopedic issues that affect standing balance control were included. All participants gave written informed consent before their participation in the experiments, and all methods were reviewed and approved by the University of Waterloo Research Ethics Board (REB #47239).

### Experimental Protocol

Before data collection, participants were briefly exposed to electrical vestibular stimulation (EVS) for 10 seconds to familiarize them with the sensation, allow them to report any discomfort, and enable the experimenter to adjust electrode placement or gel if necessary. Participants were permitted multiple familiarization exposures before data collection began.

Quiet-standing EVS trials were conducted both before and immediately after the VR exposure. Each trial lasted 120 seconds, with continuous EVS throughout. Participants were instructed to stand quietly on the force plate (model 4060-05; Bertec, Columbus, OH) with their feet shoulder-width apart, arms relaxed at their sides, and eyes fixated on a taped ‘X,’ subtending a visual angle of ∼0.76°. They were asked to remain as still as possible without locking their knees until the experimenter indicated that the trial was complete. These standardized instructions ensured consistent posture and task goals across participants.

Bipolar binaural EVS was delivered percutaneously above the mastoid processes to stimulate the nearby vestibular afferents. The anode was worn on the right and the cathode on the left. When facing forward, this configuration evokes coherent mediolateral postural sway (33,42–44). A zero mean stochastic signal was generated for each trial using a custom LabVIEW program (National Instruments Corp., Austin, TX), digitally low-pass filtered at 25 Hz (33,35,45), converted to analog, and delivered through a linear isolated stimulator (STMISOLA; Biopac Systems Inc., Goleta, CA). The EVS signal had a peak amplitude of **±** 4.5 mA and a root mean square of 1.275 mA.

After completing the quiet-standing EVS trials, participants donned the Meta Quest 1 VR system, which provides dual OLED displays (1,600 × 1,440 pixels per eye; 72 Hz refresh).

Participants stood on the force plate with continuous EVS for the entire duration of the VR trial, following the same postural instructions as during the quiet-standing EVS trials, except that they held the VR controllers at their sides. Participants were exposed to a highly nauseogenic, passive rollercoaster simulation (*Epic Roller* C*oasters*; Meta Inc.) for up to 20 minutes or until they voluntarily ended the session due to discomfort. Prior to starting, participants were informed that an FMS score above 15 served as a checkpoint to consider discontinuing the session, although the final decision was left to them. A spotter provided safety support by lightly steadying a participant at the shoulder if sway approached a potential loss of balance. The VR content was passive, meaning no user input or head movement was required to progress through the simulation. Each rollercoaster ride lasted approximately 2.5 minutes, with no breaks between successive rides.

Cybersickness was rated every 2 and a half minutes (i.e., end of each rollercoaster ride) using the Fast Motion Sickness (FMS) Scale (10), a verbal 20-point scale ranging from 0 (“no sickness”) to 20 (“severe sickness”). Following the VR exposure, participants repeated the quiet-standing EVS trial and then completed the Simulator Sickness Questionnaire (SSQ) to further quantify exposure symptoms (10,46).

### Outcome Measures

Participants were categorized into non-sick (FMS < 5), medium-sick FMS ≥ 5 who completed the task), and high-sick (participants who terminated the VR exposure early; dropout group (47)). This categorical approach captures meaningful differences in both symptom severity and functional outcomes during the VR task. Importantly, the high-sick group was defined by intolerance-driven cessation rather than peak FMS score, as some participants terminated the trial before reaching the highest ratings. Individuals in the medium-sick group completed the task but experienced noticeable symptoms, whereas non-sick participants completed it with minimal discomfort. Treating dropout as a separate outcome captures an important dimension of susceptibility that would be obscured by continuous measures. This grouping approach is further justified by the coherence and gain analyses, which pool data by concatenating time-series across individuals to generate group-level frequency-domain estimates (outlined below). Additionally, regression approaches using FMS as a continuous variable assume linear relationships and equal intervals between ratings; however, the FMS is an ordinal scale, where differences between scores (e.g., 2–4 vs. 10–12) do not necessarily represent equivalent perceptual changes in sickness.

EVS and force plate ground reaction forces and moments were sampled at 1,024 Hz using a custom LabVIEW script (Data Acquisition Board: National Instruments Corp., Austin, TX; LabVIEW: National Instruments Corp.) over a 120 second period for quiet-standing EVS trials and a 1200 second period for the VR trial. Force plate data were amplified online using an internal digital pre-amplifier and stored for off-line analysis. CoP was calculated offline using a custom MATLAB script (R2023b, MathWorks Natick, MA) from ground reaction forces and moments from the force plate. The ML-CoP was analyzed, as both EVS and VR evoke mediolateral sway. CoP is often used in balance-focused studies and provides an estimate of the neural mechanisms underlying the attempt to control the center of mass projection (1,48–51).

CoP data were low-pass filtered at 40 Hz (dual-pass, 4th-order Butterworth) to preserve the influence of the upper frequencies of EVS (≤ 25 Hz).

Baseline-corrected root mean square (RMS) of ML-CoP displacement was used to quantify sway amplitude during the pre-and post-VR quiet-standing EVS trials and the VR exposure, and how it relates to sickness severity. ML-CoP RMS was computed across the full quiet-standing EVS trials, the first and second halves of the VR trial (for participants that did not dropout), and the first minute of VR containing the initial roller-coaster drop. Data were baseline-corrected by subtracting the mean ML-CoP from each segment. Percent change was calculated between pre-and post-VR quiet-standing EVS trials and between the first and second halves of the VR game to assess sway adaptation related to sickness severity. All calculations were performed using a custom MATLAB script.

### Linear Systems Analyses

A linear systems framework quantified the relationship between EVS input and ML-CoP output in the frequency domain. Coherence and gain estimates between the EVS and ML-CoP were calculated using the NeuroSpec 2.0 code. NeuroSpec 2.0 (www.neurospec.org) is a freely available archive of MATLAB code intended for statistical signal processing and based on the methods of Halliday et al. (31). It is well established in vestibulomotor research (2,33,35,38–40,45,52,53).

Coherence quantified the proportion of ML-CoP variance explained by EVS, representing vestibulomotor coupling. It is a measure of correlation bounded between 0 and 1 for each frequency window and indicates where in the frequency spectrum signals are correlated. Zero represents the case of independence, while one represents the case of a perfect linear relationship. Calculating relative contributions to balance control can give insight into how the CNS weighs sensorimotor processes. Coherence is calculated using Eq. 1, where the cross-spectrum of the input and output is divided by the input and output auto-spectra (x is input signal, i.e., EVS; y is output signal, i.e., ML-CoP) for each frequency window (at a given time for time-frequency analyses, where time is represented by the 𝜏). For our purposes, we defined EVS as the input signal and ML-CoP as the output signal.

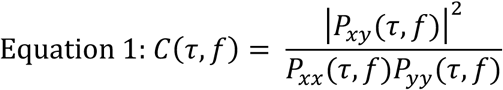

Gain reflects the scale relationship between input and output signals; in this study, it reflects the amplitude of the ML-CoP displacement to a given EVS current. Whereas coherence gives insight into the relative contributions of vestibular information, gain gives insight into how large the total response to EVS is. Interpreting gain is conditional on significant coherence between signals, so any frequencies where coherence was not significant were excluded from the gain analyses. It is calculated by using Eq. 2, where the magnitude of the cross-spectrum between input and output is divided by the input auto-spectrum for each frequency window (at a given time for time-frequency analyses, where time is represented by the 𝜏).

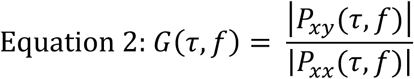

For sickness group-level analyses, participant data were concatenated into arrays to produce group coherence and gain. Coherence and gain were computed from 0.5–30 Hz using 2-second non-overlapping windows (2,048 samples) beginning at EVS onset. A 2 second window (0.5 Hz resolution) was needed to resolve frequencies at 0.5 Hz, where most CoP power occurs (54). The frequency range was selected because vestibulomotor coherence has been found up to 30 Hz in previous studies (2,33,35,38,45,52).

Concatenation was performed separately for the pre-and post-VR quiet-standing EVS trials and for the first and second halves of the VR trial. Data were concatenated into arrays such that input and output data were matched sequentially (i.e., EVS data from participant 1 aligned with ML-CoP data from participant 1). For non-sick, this approach yielded 720 two-second windows for the pre-and post-VR quiet-standing EVS trials (n = 12) and 3,600 windows for each half of the VR trials. For medium-and high-sick, this approach yielded 780 two-second windows for the pre-and post-VR quiet-standing EVS trials (n = 13) and 3,900 windows for each half of the VR trials.

Within-subject mean coherence across 0.5–15 Hz was also computed to assess inter-individual consistency. This frequency range was chosen during post hoc analyses based on the quiet-standing EVS trials, since there was significant coherence up to 15 Hz. Mean coherence was calculated for each participant during the pre-and post-VR quiet-standing EVS trials, as well as the first and second halves of the VR trial separately.

During the VR trial, individual coherence spectra were also computed from 0.5–30 Hz using 2-second windows across consecutive 2.5-minute segments (8 total). The mean EVS-ML-CoP coherence for each participant was then calculated across 0.5–15 Hz within each segment. To assess group-level trends, mean coherence values at each time point were averaged across participants in the non-sick and medium-sick groups. The same analysis was performed for the high-sick group for up to four segments due to truncated VR exposure, with participants requiring balance support excluded. This approach captured changes in vestibulomotor coupling over the course of the VR exposure. Gain was not analyzed in this way because it is not interpretable at frequencies where coherence is insignificant, and coherence differences were more pronounced, making temporal changes easier to detect.

## Statistical Analyses

All analyses were performed in MATLAB (R2023b). Normality was assessed using the Shapiro-Wilk test (*p* < 0.05 indicated non-normality). Effect sizes were reported as η² (eta-squared) for ANOVAs, non-parametric η² for Kruskal-Wallis tests, Cohen’s *d* for independent-samples t-tests, and rank-biserial correlation (𝑟_$%_) for Mann-Whitney U tests and Wilcoxon signed-rank tests.

To ensure that sickness groups were statistically different in symptoms reported with the SSQ, one-way ANOVAs (or Kruskal-Wallis tests) was conducted between non-sick, medium-sick and high-sick groups. A simple regression was also calculated between peak FMS score and total SSQ score. To determine whether specific SSQ symptom domains (nausea, oculomotor, or disorientation) contributed disproportionately to sickness severity, simple regressions were performed between each subscale total and peak FMS score.

One-way ANOVAs (or Kruskal-Wallis tests) compared ML-CoP RMS across sickness groups (non-, medium-, high-sick) for pre-VR quiet-standing EVS trials, post-VR quiet-standing EVS trials, percent change from pre-to post-VR quiet-standing EVS trials, percent change from first to second VR halves, and first minute of the roller-coaster drop. Linear regressions were calculated between peak FMS score and these same ML-CoP metrics to see if FMS scores were sensitive beyond group classifications.

Significant EVS-ML-CoP coherence was confirmed using a mean 95% confidence limit of coherence which was calculated across all windows for individual and concatenated data.

Significant coherence served as a prerequisite for gain analyses. The frequency ranges with significant coherence between concatenated sickness conditions were recorded and compared.

Differences in coherence within sickness groups from pre-to post-VR quiet-standing EVS trials, as well as between sickness groups for pre-and post-VR trials, were examined at the group level using difference-of-coherence tests based on the methods of Rosenberg et al. (32) and Amjad et al. (55). This statistic is a modified *χ*^2^ test, testing the assumption that the coherence estimates are equal with normally distributed variance across conditions. Standardized coherence differences were calculated across 0-15 Hz between pre-and post-VR quiet-standing EVS trials within each sickness group and across groups. Similar comparisons were performed between the first and second halves of VR exposure within groups (non-sick and medium-sick only). Ninety-five percent confidence limits were derived using the Fisher transform (tanh⁻¹) of the square root of the coherence values. Frequencies at which standardized coherence differences exceeded the 95% confidence limits were considered statistically significant.

Differences in coherence were also examined using mean coherence values across participants to better capture inter-individual variability and effect size. Mean EVS-ML-CoP coherence across 0.5–15 Hz was compared within and between groups using the appropriate statistical test for the pre-and post-VR quiet-standing EVS trials and the first and second halves of the VR exposure.

For the VR trials, temporal changes in group-level coherence were analyzed using linear regression of mean coherence to each time segment. For individual-level analyses, y-intercepts, slopes, and R² values describing coherence over time were computed for each participant.

Independent t-tests were then used to compare these metrics between the non-sick and medium-sick groups, allowing assessment of inter-individual variability and group differences in coherence trajectories.

Significant differences in EVS-ML-CoP gain between pre-and post-VR quiet-standing EVS trials and between sickness groups pre-and post-VR were calculated using pointwise 95% confidence limits for all windows of group concatenated data across conditions. The same analysis was conducted within groups for the first and second halves of the VR exposure.

Regions of nonoverlapping confidence limits were distinguished when significant differences occurred (2,33). When overlapping between the confidence limits occurred, gain values were not considered significantly different. Within the regions where gain was significantly different across conditions, the mean and standard deviation of percentage change were recorded to capture effect size.

## Results

### Post Collection Exclusion

Spotter assistance was required for three participants, all in the high-sickness group. In these cases, the spotter lightly stabilized the participants’ shoulders to prevent loss of balance. Due to variable dropout times, group-level EVS-ML-CoP coherence analyses were not conducted for high-sickness VR trials; however, data from these participants were included for pre-and post-VR quiet-standing EVS analyses.

### Sickness

Participants exhibited a wide range of cybersickness responses during the VR rollercoaster task, enabling classification into non-sick and medium-sick groups based on peak FMS scores, and a high-sick group defined by early termination of the VR exposure. Non-sick (peak FMS < 5) had 12 participants (Fig. 1A), medium-sick (peak FMS ≥ 5 without VR dropout) had 13 (Fig.1B), and high-sick (VR discontinuation before 20 minutes) had 13 (Fig. 1C).

**Figure 1.**
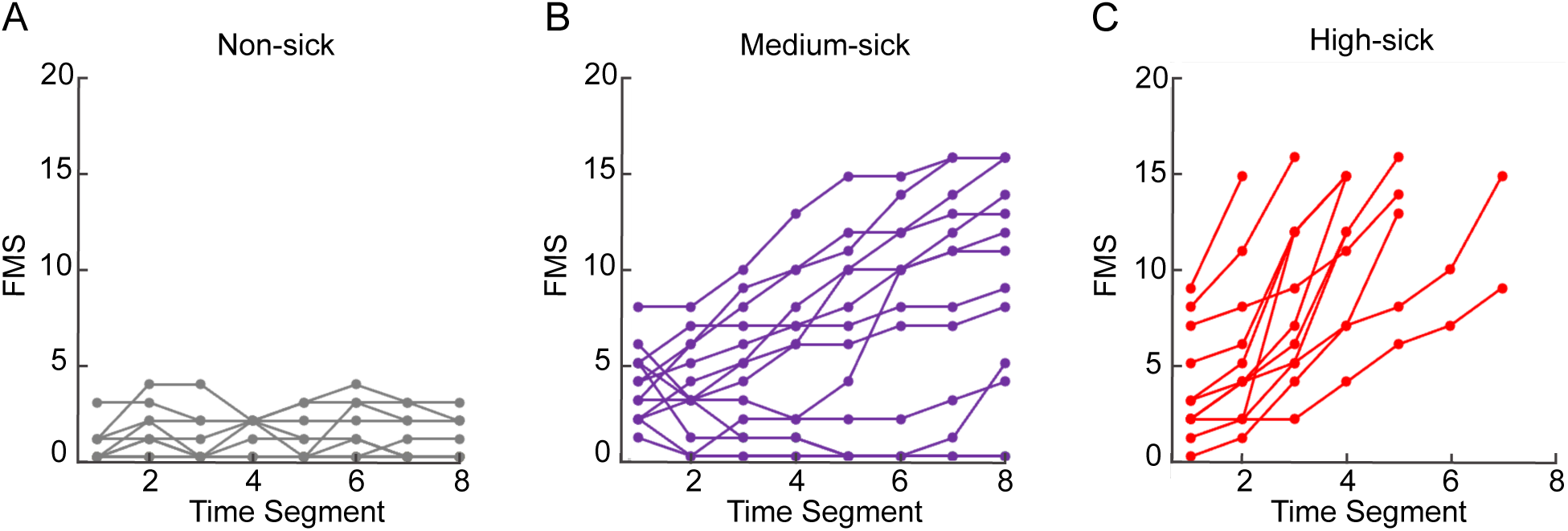
Individual FMS scores over time during VR exposure. (A–C) FMS scores for non-sick (A), medium-sick (B), and high-sick (C) participants, plotted across consecutive 2.5-minute time segments of the VR trial. Each line represents an individual participant.

A one-way ANOVA revealed a significant main effect of sickness group on SSQ scores, with a large effect size (*p* < 0.001, η² = 0.489; Fig. 2). Post hoc Tukey tests indicated that SSQ scores in the non-sick group (17.75 ± 3.35) were significantly lower than those in both the medium-sick (75.03 ± 15.10; *p* = 0.002) and high-sick groups (105.66 ± 9.62; *p*< 0.001), whereas the medium-and high-sick groups did not differ significantly.

**Figure 2.**
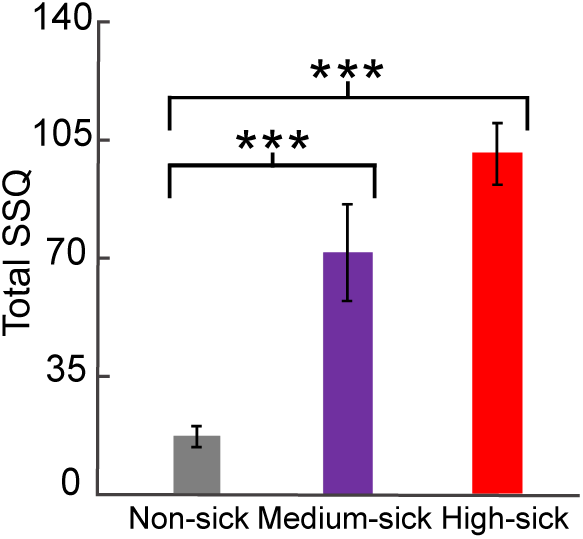
Total SSQ scores for non-sick, medium-sick, and high-sick groups. Error bars represent ±1 standard error of the mean (SEM). Asterisks indicate significant pairwise differences (* *p* < 0.05, ** *p* < 0.01, *** *p* < 0.001).

A simple linear regression revealed a strong linear relationship between FMS and SSQ scores (*p* < 0.001, R² = 0.58; Fig. 3A). Among SSQ subscales, nausea showed the strongest association with peak FMS scores (*p* < 0.001, R² = 0.67; Fig. 3B), exceeding associations with oculomotor (*p* < 0.001, R² = 0.42; Fig. 3C) and disorientation sub scores (*p* < 0.001, R² = 0.48; Fig. 3D).

**Figure 3.**
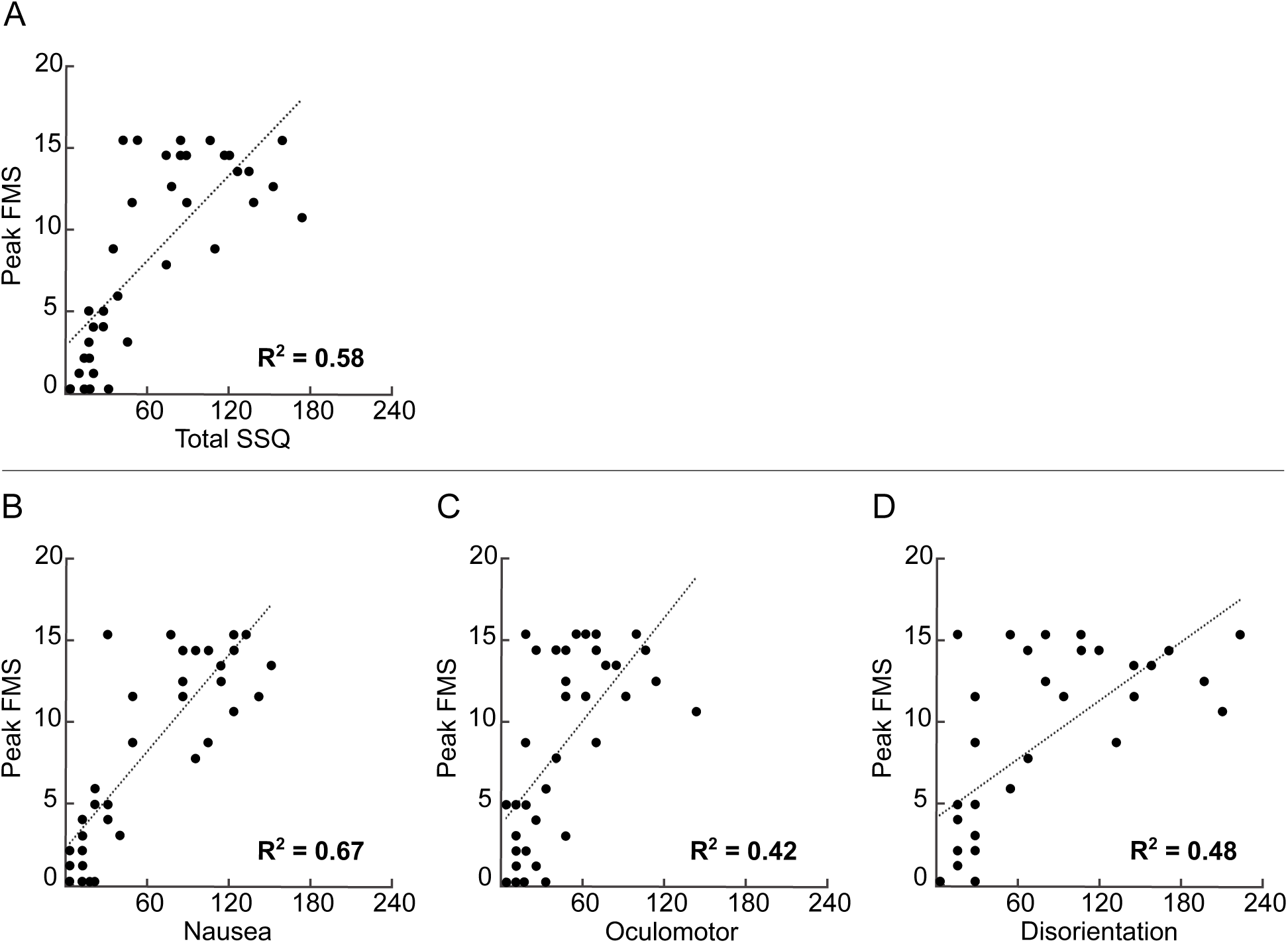
Relationship between peak FMS scores and SSQ measures across participants. (A–D) Peak FMS scores plotted against total SSQ scores (A), nausea (B), oculomotor (C), and disorientation (D) subscale scores. Each point represents an individual participant, and dotted lines indicate best-fit linear regressions. R² values where *p* <0.05 are in bold.

### ML-CoP RMS

To determine whether cybersickness was accompanied by changes in postural stability, we first examined ML-CoP RMS across conditions and sickness groups. During the pre-VR quiet-standing EVS trials, a Kruskal-Wallis test revealed no significant differences in ML-CoP RMS across sickness groups (non-sick: 3.99 × 10⁻³ ± 6.17 × 10⁻⁴; medium-sick: 4.40 × 10⁻³ ± 8.63 × 10⁻⁴; high-sick: 3.51 × 10⁻³ ± 4.67 × 10⁻⁴; Fig. 4A).

**Figure 4.**
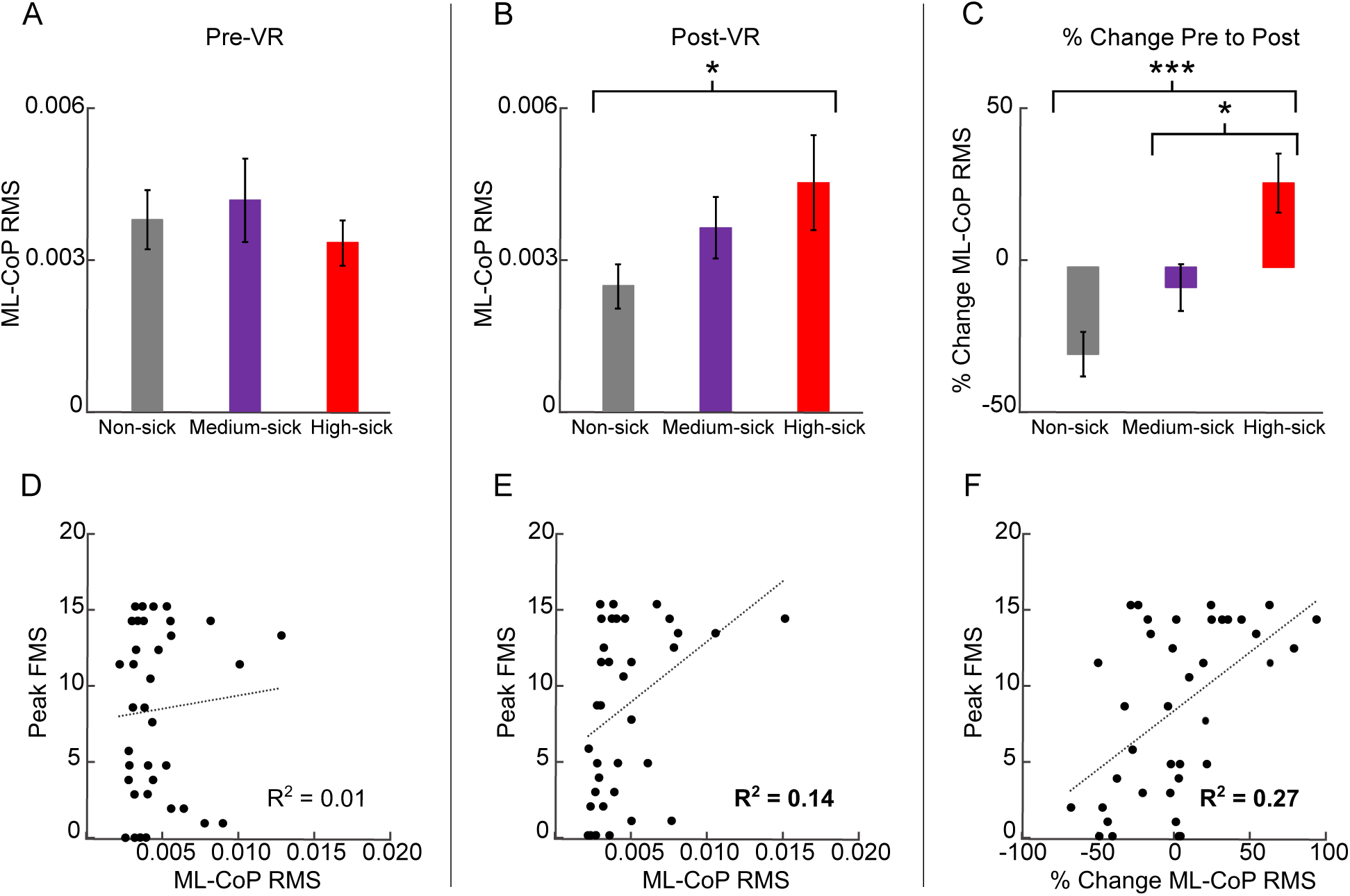
ML-CoP RMS and its relationship with peak FMS scores during quiet-standing EVS trials. (A–C) ML-CoP RMS measured prior to VR exposure (A), following VR exposure (B), and percent change from pre-to post-VR exposure (C) across non-sick, medium-sick, and high-sick participants. Bars represent group means, with error bars indicating ± SEM. Asterisks indicate significant pairwise differences (* *p* < 0.05, ** *p* < 0.01, *** *p* < 0.001). (D–F) Peak FMS scores plotted against ML-CoP RMS during pre-VR trials (D), post-VR trials (E), and percent change from pre-to post-VR (F). Each point represents an individual participant, and dotted lines indicate best-fit linear regressions. R² values where *p* <0.05 are in bold.

In contrast, post-VR ML-CoP RMS differed significantly across groups (*p* = 0.043, η² = 0.123; Fig. 4B). Post hoc Tukey tests indicated that ML-CoP RMS was greater in the high-sick group than in the non-sick group (*p* = 0.043), with no other group differences observed. ML-CoP RMS increased with sickness severity (non-sick: 2.60 × 10⁻³ ± 4.62 × 10⁻⁴; medium-sick: 3.82 × 10⁻³ ± 6.39 × 10⁻⁴; high-sick: 4.76 × 10⁻³ ± 9.87 × 10⁻⁴).

A one-way ANOVA revealed a significant effect of sickness group on the change in ML-CoP RMS from pre-to post-VR (*p* < 0.001, η² = 0.392; Fig. 4C). Post hoc Tukey tests indicated that the high-sick group exhibited greater increases in ML-CoP RMS than both the non-sick (*p* < 0.001) and medium-sick groups (*p* = 0.016), whereas the non-sick and medium-sick groups did not differ. Changes in ML-CoP RMS were −30.06 ± 7.67% (non-sick), −7.00 ± 8.19% (medium-sick), and 29.18 ± 10.30% (high-sick).

Simple linear regressions revealed patterns consistent with the group comparisons. The relationship between peak FMS score and ML-CoP RMS during pre-VR quiet-standing EVS trials was not significant, whereas a significant association was observed for post-VR ML-CoP RMS (*p* = 0.022, R² = 0.14; Fig. 4E). The relationship was also significant when examining the percent change in ML-CoP RMS from pre-to post-VR (*p* < 0.001, R² = 0.27; Fig. 4F).

When examining the first and second halves of the VR trial, as well as the percent change between halves for the non-sick and medium-sick groups, trends similar to those observed during the quiet-standing EVS trials were evident, however, no comparisons reached statistical significance using Mann-Whitney U tests or independent-samples *t*-tests. ML-CoP RMS during the first half was 8.29 × 10⁻³ ± 1.55 × 10⁻³ for the non-sick group and 8.37 × 10⁻³ ± 9.38 × 10⁻⁴ for the medium-sick group (Fig. 5A). During the second half, ML-CoP RMS was 7.78 × 10⁻³ ± 1.59 × 10⁻³ for the non-sick group and 1.01 × 10⁻² ± 1.55 × 10⁻³ for the medium-sick group (Fig. 5B). The percent change in ML-CoP RMS from the first to the second half was −2.74 ± 11.30% for the non-sick group and 18.74 ± 10.18% for the medium-sick group (Fig. 5C).

**Figure 5.**
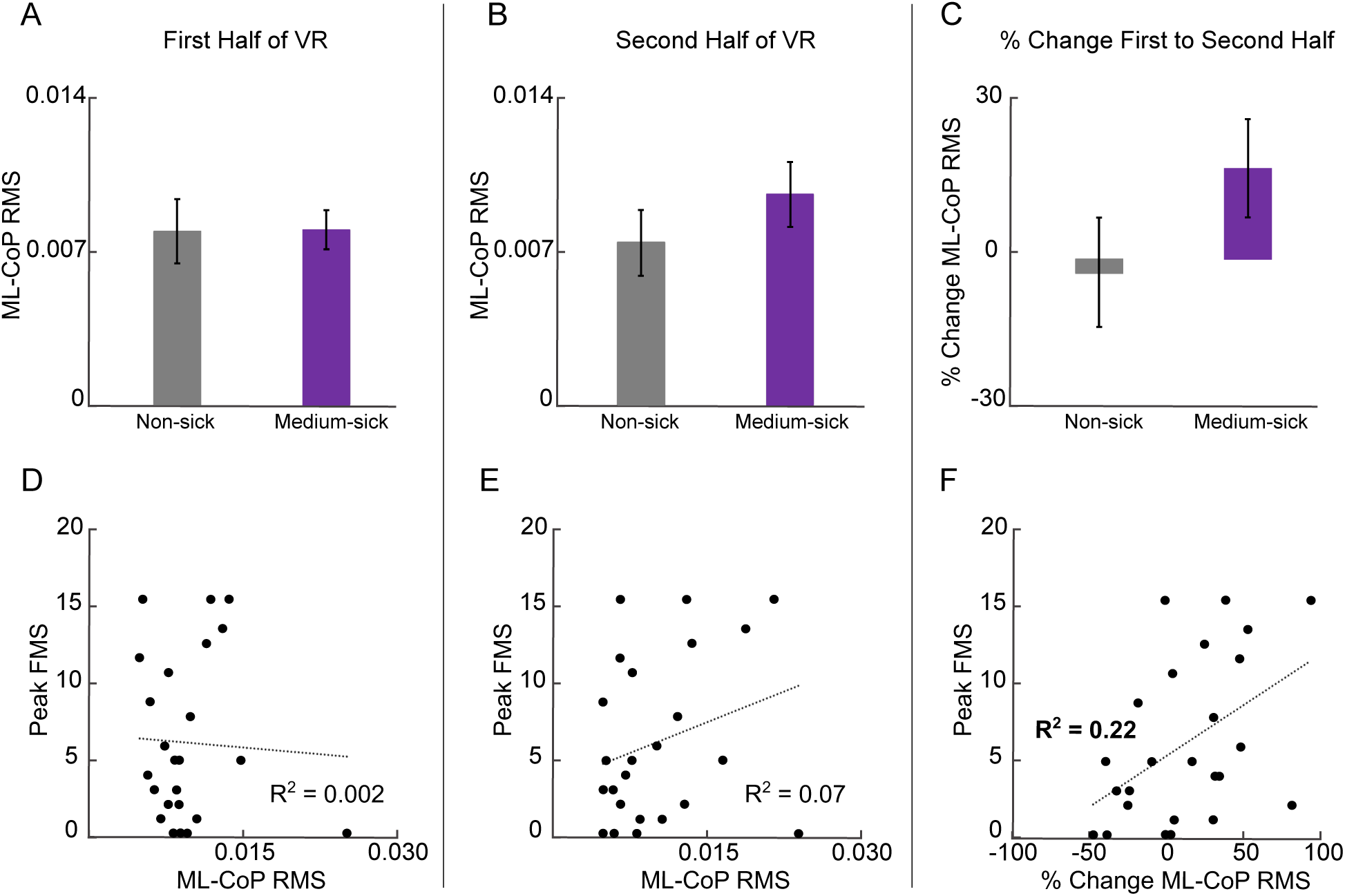
ML-CoP RMS during the VR trial and its relationship with peak FMS scores. (A–C) ML-CoP RMS averaged across the first half of the VR exposure (A), second half (B), and percent change between halves (C) for non-sick and medium-sick participants. Bars represent group means, and error bars indicate ± SEM. (D–F) Peak FMS scores plotted against ML-CoP RMS during the first half (D), second half (E), and percent change between halves (F). Each point represents an individual participant, and dotted lines indicate best-fit linear regressions. R² values where *p* <0.05 are in bold.

Simple linear regressions revealed patterns consistent with the group comparisons, although statistical significance was observed for the percent change from the first to the second half of the VR trial. The relationships between peak FMS score and ML-CoP RMS during the first and second halves of VR were not significant (first half: R² = 0.002; second half: R² = 0.072; Fig. 5D, E). In contrast, the association between peak FMS score and the percent change in ML-CoP RMS from the first to the second half was significant (*p* = 0.019, R² = 0.22; Fig. 5F).

Transient responses to salient visual perturbations were examined to assess whether early instability predicted sickness severity. When examining ML-CoP RMS during the first minute of the VR trial, which included the initial rollercoaster drop, a one-way ANOVA revealed a significant main effect of sickness group (*p* < 0.001, η² = 0.329; Fig. 6A). Post hoc Tukey tests indicated that ML-CoP RMS did not differ between the non-sick and medium-sick groups but was significantly greater in the high-sick group than in both the non-sick (*p* = 0.004) and medium-sick groups (*p* = 0.003). Mean ML-CoP RMS values were 6.93 × 10⁻³ ± 9.24 × 10⁻⁴ (non-sick), 6.78 × 10⁻³ ± 1.16 × 10⁻³ (medium-sick), and 1.52 × 10⁻² ± 2.70 × 10⁻³ (high-sick). Additionally, linear regression revealed a significant positive association between peak FMS score and ML-CoP RMS during this first minute (R² = 0.11, *p* = 0.046; Fig. 6B).

**Figure 6.**
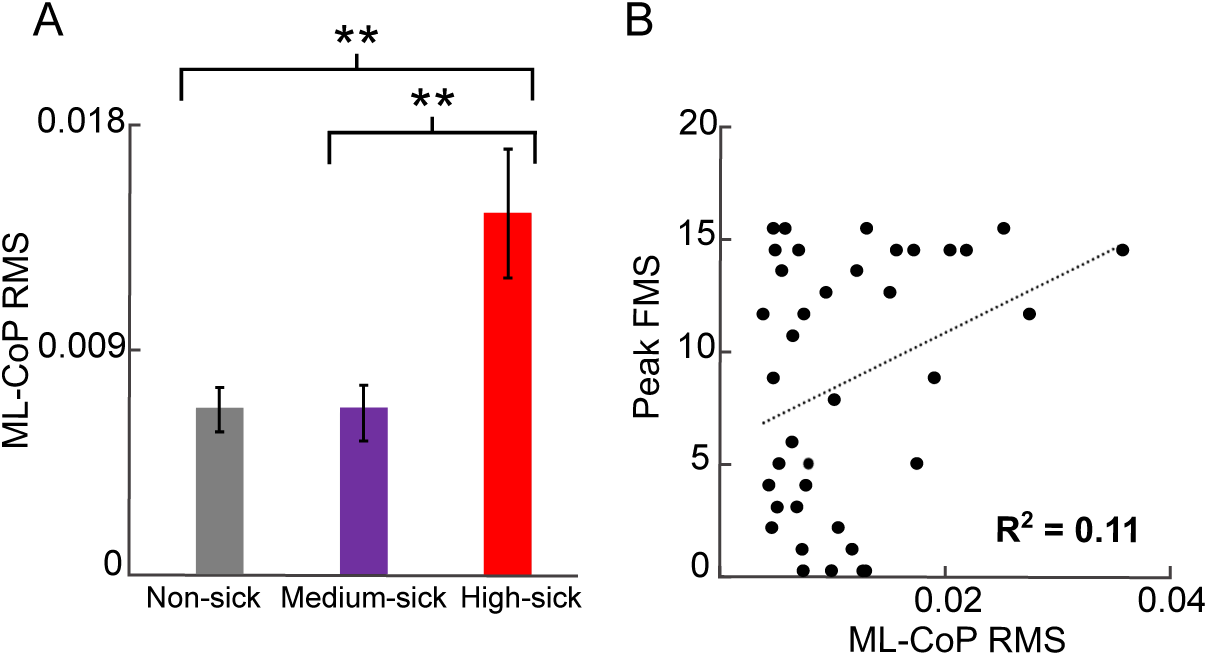
ML-CoP RMS during the first minute of the VR trial, including the initial rollercoaster drop. (A) Mean ML-CoP RMS for non-sick, medium-sick, and high-sick participants. Bars represent group means and error bars indicate ± SEM. Asterisks indicate significant pairwise differences (* *p* < 0.05, ** *p* < 0.01, *** *p* < 0.001). (B) Peak FMS scores plotted against ML-CoP RMS during the first minute of the VR trial. Each point represents an individual participant, and the dotted line indicates the best-fit linear regression. R² values where *p* <0.05 are in bold.

### Coherence

To determine whether differences in postural behavior were associated with vestibulomotor weighting, we next examined EVS-ML-CoP coherence. Significant concatenated group-level coherence was present across all sickness groups, although the frequency ranges varied by group and condition. In the non-sick group, coherence was observed during both pre-and post-VR quiet-standing EVS trials across 0.5–9.5 Hz (Fig. 7A). In the medium-sick group, no significant coherence was detected pre-VR, whereas post-VR coherence emerged at 0.5–2 Hz and 3.5–5 Hz (Fig. 7B). In the high-sick group, coherence was present pre-VR across 1–10.5 Hz, 11–13 Hz, and 14–15.5 Hz, and post-VR across 0.5–9 Hz (Fig. 7C). The robustness of these group-level patterns was generally supported by individual participant spectra (Fig. 7D–F).

**Figure 7.**
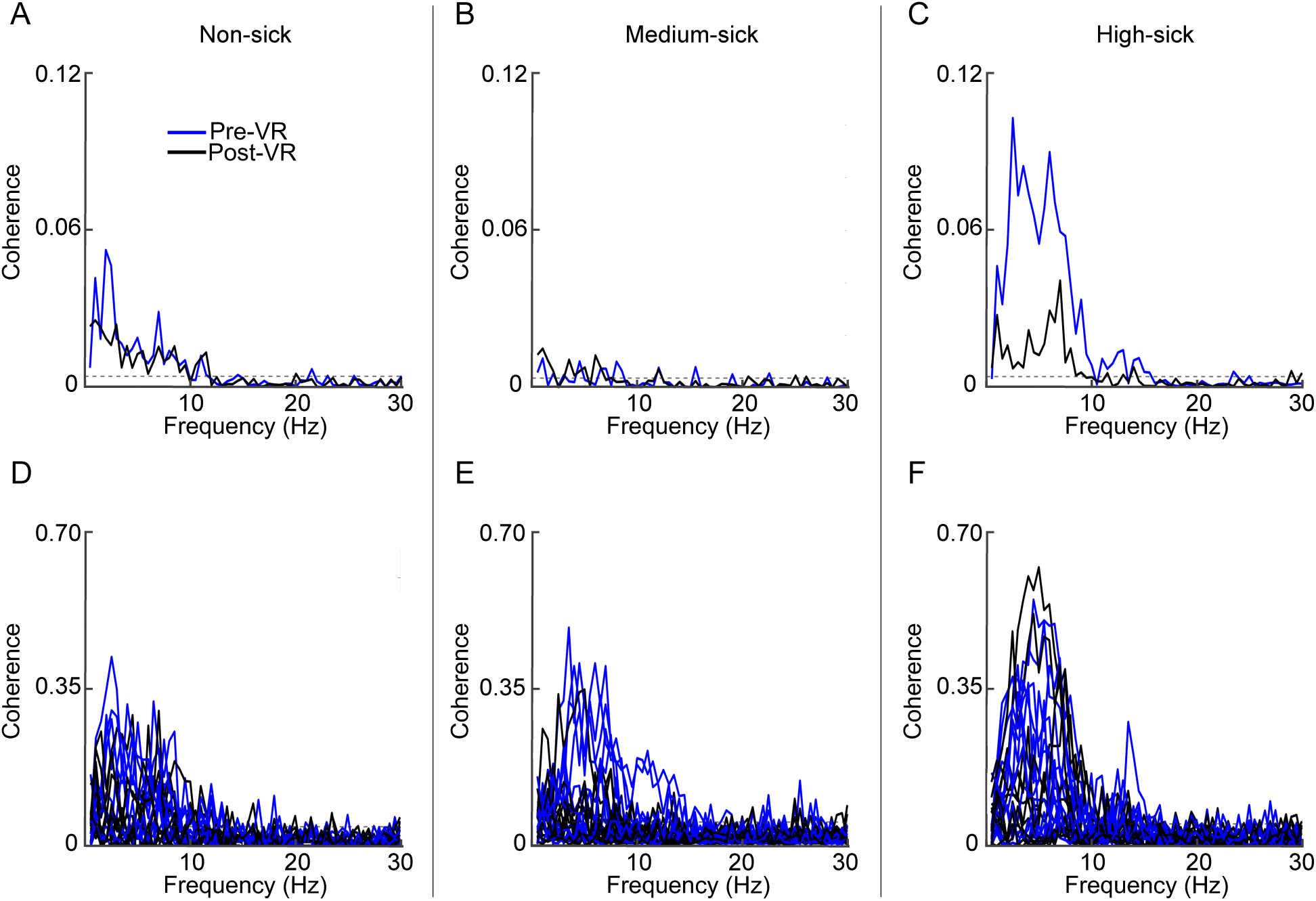
EVS-ML-CoP coherence spectra during quiet-standing EVS trials for each sickness group. (A–C) Group-level coherence calculated from concatenated data for the non-sick (A), medium-sick (B), and high-sick (C) groups, shown separately for pre-VR (blue) and post-VR (black) trials. (D–F) Individual participant coherence spectra for the non-sick (D), medium-sick (E), and high-sick (F) groups, with each line representing a single participant and colors indicating pre-VR (blue) and post-VR (black) conditions. Coherence is plotted as a function of frequency. A dashed horizontal line indicates the 95% confidence limit threshold for significant coherence.

We next assessed whether vestibulomotor coupling changed following VR exposure.

Significant differences in concatenated group-level coherence between pre-and post-VR quiet-standing EVS trials were observed only in the high-sick group. In this group, pre-VR coherence was greater than post-VR coherence across the 1.5–6.5 Hz and 7.5–9 Hz frequency ranges (Fig. 8C). No significant pre-post differences in concatenated coherence were detected in the non-sick or medium-sick groups (Fig. 8A, B).

**Figure 8.**
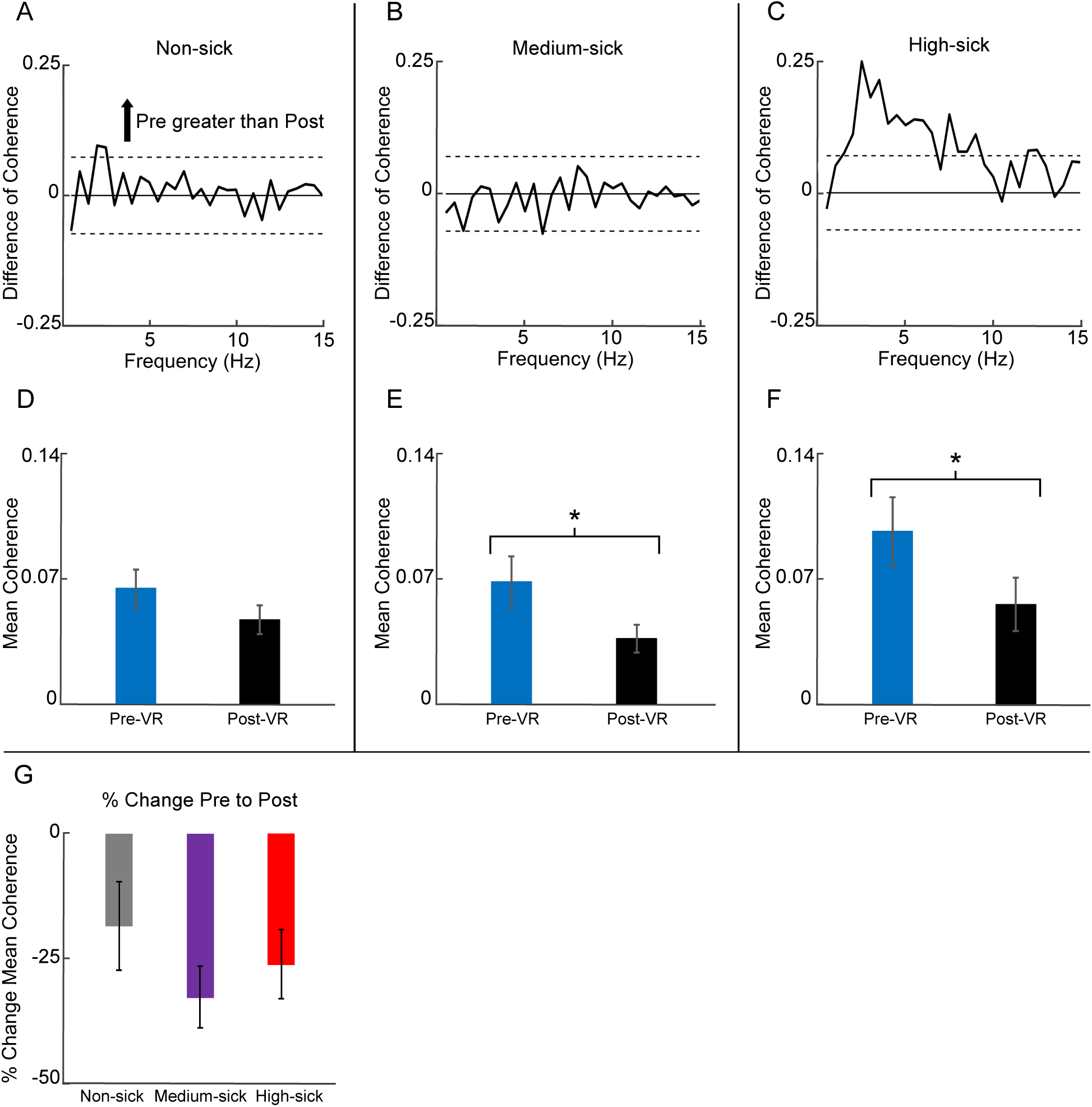
Difference-of-coherence and average coherence results for quiet-standing EVS trials across sickness groups. (A–C) Difference-of-coherence spectra comparing pre-and post-VR trials for the non-sick (A), medium-sick (B), and high-sick (C) groups, plotted as a function of frequency. Horizontal lines indicate the 95% confidence limits used for the difference-of-coherence test. (D–F) Mean EVS-ML-CoP coherence averaged across frequencies (0–15 Hz) for pre-VR (blue) and post-VR (black) trials in the non-sick (D), medium-sick (E), and high-sick (F) groups. Bars represent group means and error bars indicate ± SEM. Asterisks indicate significant pairwise differences (* *p* < 0.05, ** *p* < 0.01, *** *p* < 0.001). (G) Percent change in mean EVS-ML-CoP coherence from pre-to post-VR trials across sickness groups. Bars represent group means and error bars indicate ± SEM.

When coherence was averaged across frequencies from 0–15 Hz and across participants, non-sick participants did not show a significant change from pre-to post-VR (pre: 0.069 ± 0.011; post: 0.050 ± 0.0087; Fig. 8D). Medium-sick participants exhibited elevated pre-VR coherence (0.073 ± 0.016) that decreased significantly following VR exposure (post: 0.039 ± 0.008; Wilcoxon signed-rank test, *p* = 0.011, 𝑟_rb_ = 0.678; Fig. 8E). Similarly, high-sick participants showed the greatest pre-VR coherence (0.103 ± 0.020), which declined significantly post-VR (0.060 ± 0.016; Wilcoxon signed-rank test, *p* = 0.040, 𝑟_rb_ = 0.562; Fig. 8F). No significant differences in the percent change in mean coherence from pre-to post-VR were observed across sickness groups (Kruskal-Wallis test; non-sick: −19.43 ± 9.42%; medium-sick: −34.67 ± 6.59%; high-sick: −27.62 ± 7.35%; Fig. 8G).

To directly compare baseline vestibular weighting across groups, between-group coherence differences were examined. Significant differences in concatenated coherence between the high-sick and non-sick groups were observed during pre-VR quiet-standing EVS trials, but not during post-VR trials. For these between-group comparisons, an equal number of participants was required; therefore, one participant was removed from the high-sick group to match the non-sick group. In the pre-VR condition, coherence in the high-sick group exceeded that of the non-sick group across 2.5–8 Hz (Fig. 9A). When coherence was averaged across frequencies from 0–15 Hz and across participants to account for inter-individual variability (with no participants removed), Kruskal-Wallis tests revealed no significant differences across sickness groups in either the pre-or post-VR quiet-standing EVS trials (Fig. 9C, D).

**Figure 9.**
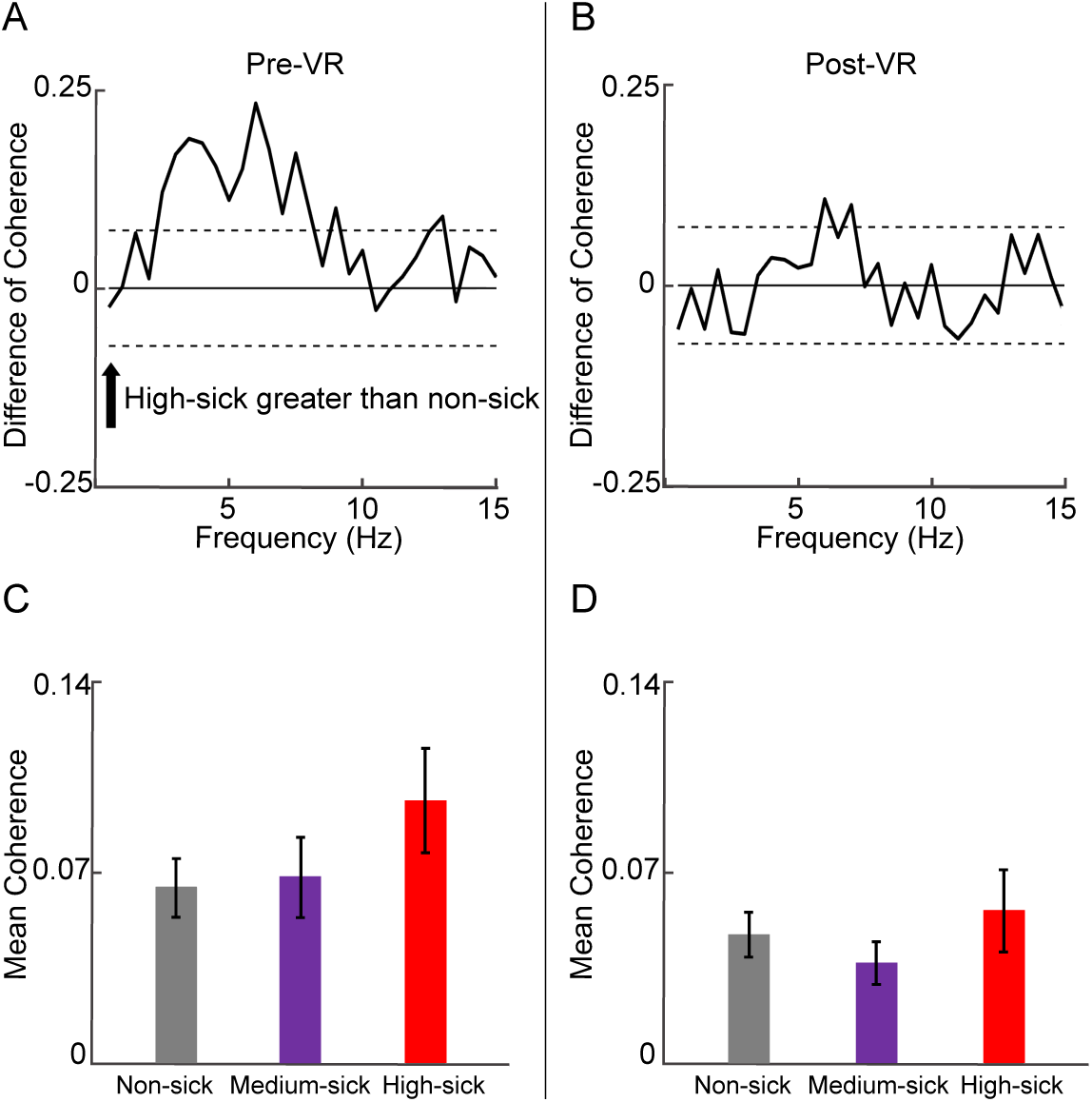
Difference-of-coherence and mean coherence comparisons between high-sick and non-sick groups during quiet-standing EVS trials. (A, B) Difference-of-coherence spectra for pre-VR (A) and post-VR (B) conditions, plotted as a function of frequency. Horizontal lines indicate the 95% confidence limits used for the difference-of-coherence tests. (C, D) Mean EVS-ML-CoP coherence averaged across frequencies (0–15 Hz) for pre-VR (C) and post-VR (D) trials across sickness groups. Bars represent group means, and error bars indicate ± SEM.

To examine how vestibulomotor coupling evolved during VR exposure, coherence was analyzed across the first and second halves of the trial. Patterns similar to those observed during the quiet-standing EVS trials were evident. In the non-sick group, coherence was present during the first half of the VR trial across 0.5–11 Hz and during the second half across 0.5–13 Hz (Fig. 10A). In the medium-sick group, coherence during the first half spanned 0.5–13 Hz, whereas during the second half it was present across 0.5–5.5 Hz, 6.5–8.5 Hz, and 9.5–11.5 Hz (Fig. 10B).

**Figure 10.**
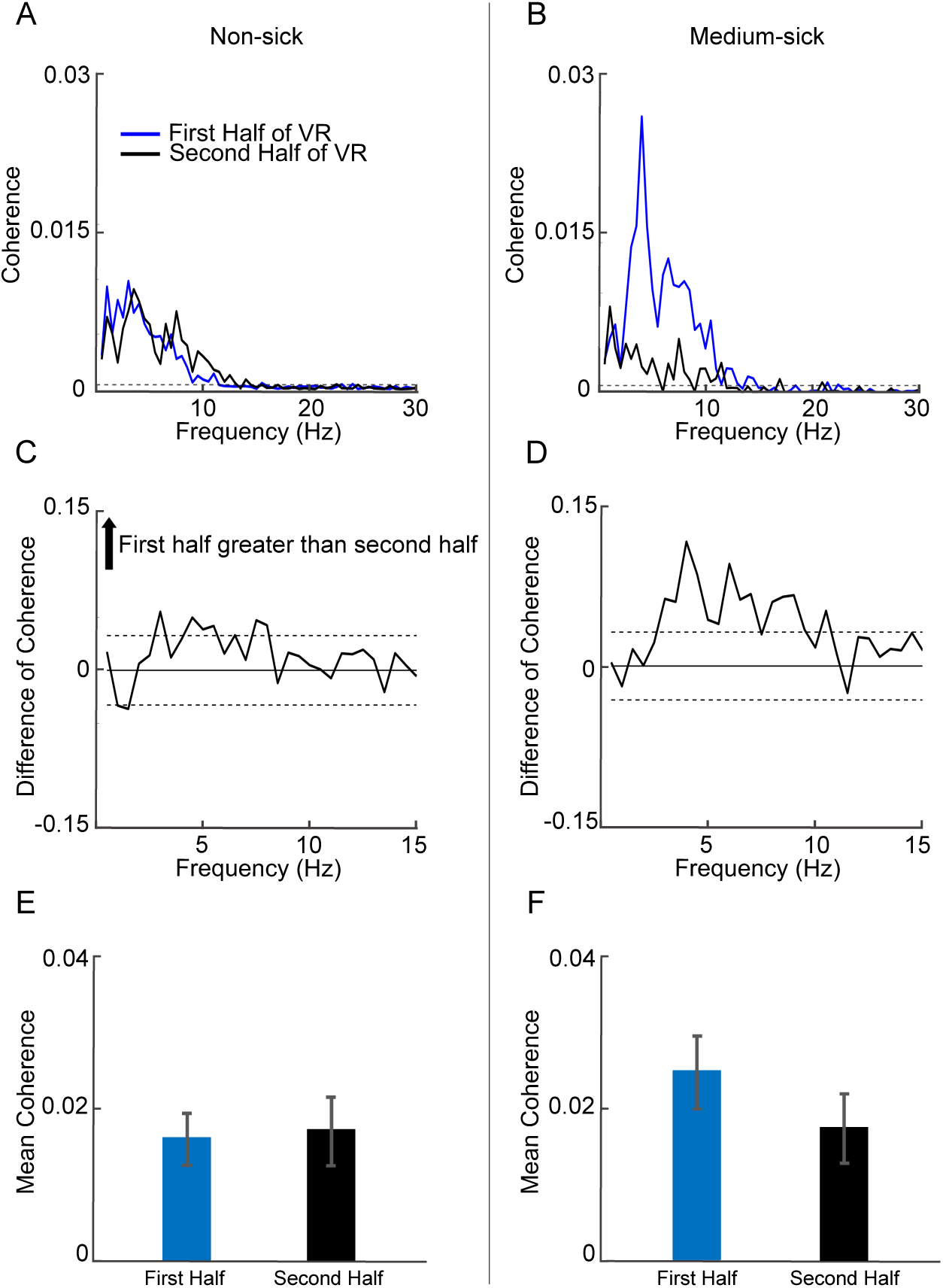
EVS-ML-CoP coherence during the VR trial for non-sick and medium-sick groups. (A, B) Concatenated group-level coherence spectra for the non-sick (A) and medium-sick (B) groups, shown separately for the first half (blue) and second half (black) of the VR exposure. Coherence is plotted as a function of frequency. A dashed horizontal line indicates the 95% confidence limit threshold for significant coherence. (C, D) Difference-of-coherence spectra comparing the first and second halves of the VR trial for the non-sick (C) and medium-sick (D) groups, plotted as a function of frequency. Horizontal lines indicate the 95% confidence limits used for the difference-of-coherence tests. (E, F) Mean EVS-ML-CoP coherence averaged across frequencies (0–15 Hz) for the first and second halves of the VR trial in the non-sick (E) and medium-sick (F) groups. Bars represent group means and error bars indicate ± SEM.

Concatenated group-level comparisons revealed differences between the first and second halves of the VR trial in both sickness groups, with more extensive differences in the medium-sick group. In the non-sick group, coherence during the first half exceeded that of the second half across 4.5–5.5 Hz (Fig. 10C). In the medium-sick group, first-half coherence exceeded second-half coherence across the 3–7 Hz and 8–9.5 Hz frequency bands (Fig. 10D).

However, when coherence was averaged across frequencies from 0–15 Hz and across participants to account for inter-individual variability, Wilcoxon signed-rank tests revealed no significant differences between halves in either sickness group, despite trends toward greater first-half coherence in the medium-sick group (non-sick: first half = 0.017 ± 0.004, second half = 0.019 ± 0.005; medium-sick: first half = 0.027 ± 0.005, second half = 0.019 ± 0.005; Fig. 10E, F).

To quantify continuous changes in vestibulomotor coupling over time, linear regressions were applied to coherence across consecutive VR segments. Linear regressions were significant for the medium-sick group but not for the non-sick group. For the medium-sick group, mean EVS-ML-CoP coherence averaged across participants for each 2.5-minute segment (eight segments total) showed a y-intercept of 0.0493, a slope of −0.0027, and a strong fit (*p* < 0.001, R² = 0.88; Fig. 11B). In contrast, the non-sick group exhibited a y-intercept of 0.0315, a shallow negative slope (−0.0005), and a weaker fit (R² = 0.21; Fig. 11A). The high-sick group included nine participants with variable dropout times across the first four segments, with some excluded due to balance assistance requirements. In this group, coherence showed a y-intercept of 0.0437, a slope of −0.0060, and a strong fit (R² = 0.97; Fig. 11C). The robustness of these group-level trends was supported by individual participant mean coherence trajectories (Fig. 11D–F).

**Figure 11.**
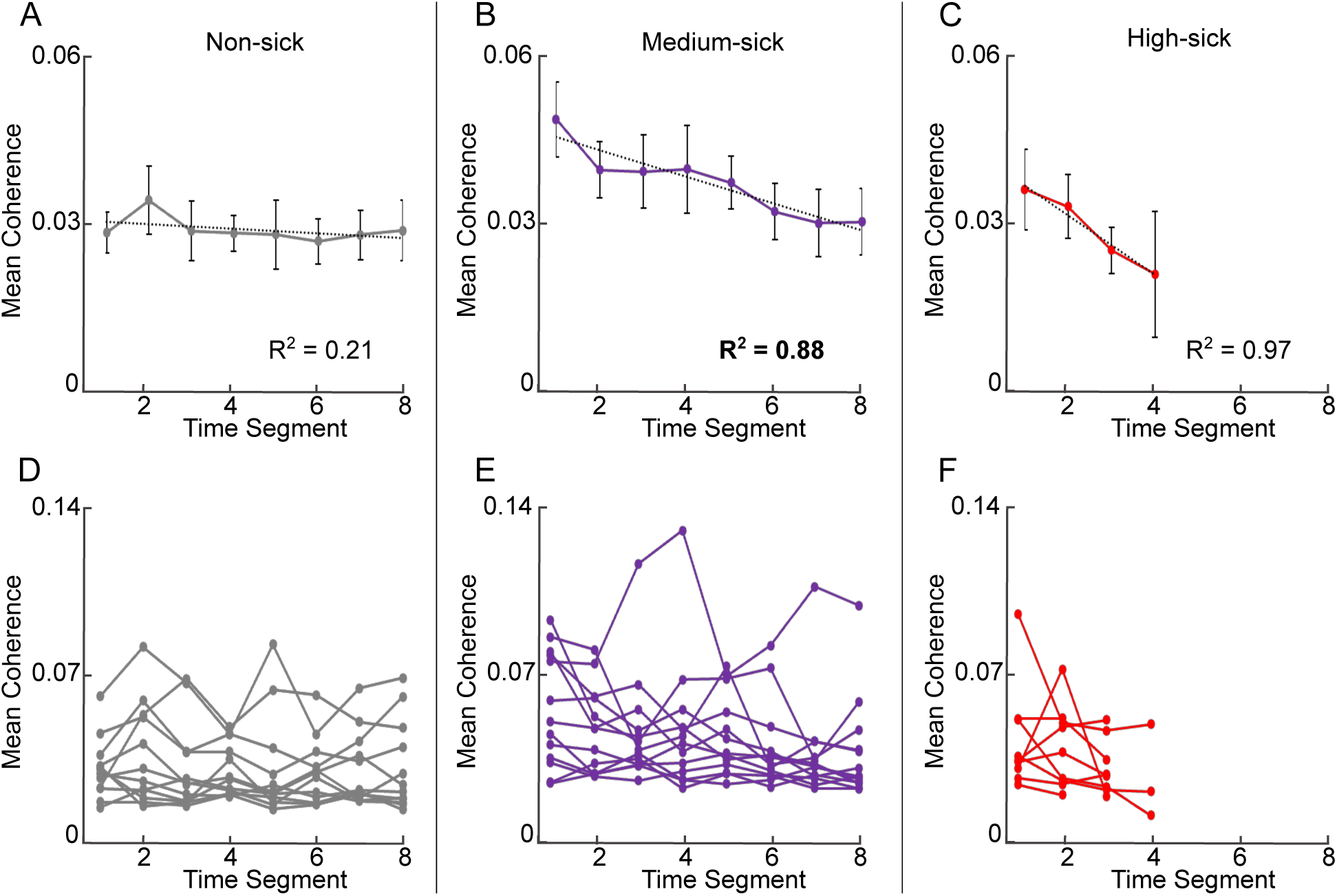
Temporal changes in EVS-ML-CoP coherence during the VR trial across sickness groups. (A–C) Group-averaged coherence plotted across consecutive 2.5-minute time segments for the non-sick (A), medium-sick (B), and high-sick (C) groups. Dotted lines indicate linear regression fits to the group-averaged data. R² values where *p* <0.05 are in bold. Error bars represent ± SEM. (D–F) Individual participant coherence values plotted across the same time segments for the non-sick (D), medium-sick (E), and high-sick (F) groups, with each line representing a single participant.

Independent-samples *t*-tests were used to assess inter-individual differences in coherence regression parameters (y-intercepts, slopes, and R² values) between the non-sick and medium-sick groups. Y-intercepts approached significance (*p*= 0.050; Fig. 12A), whereas significant group differences were observed for slopes (*p* = 0.017, Cohen’s *d* = 1.03; Fig. 12B) and R² values (*p* = 0.040, Cohen’s *d* = −0.87; Fig. 13C).

**Figure 12.**
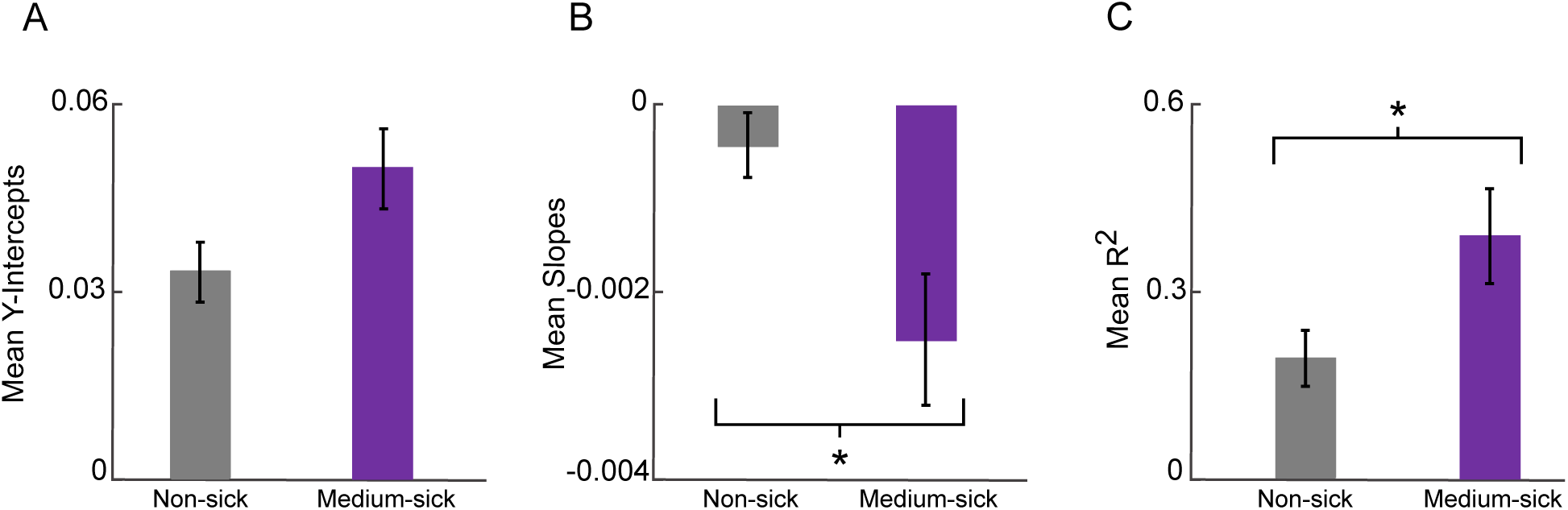
Comparison of linear regression parameters describing temporal changes in EVS-ML-CoP coherence between non-sick and medium-sick groups. (A–C) Mean y-intercepts (A), mean slopes (B), and mean R² values (C) derived from individual participant regressions of coherence across consecutive 2.5-minute VR time segments. Bars represent group means and error bars indicate ± SEM. Asterisks indicate significant pairwise differences (* *p* < 0.05, ** *p* < 0.01, *** *p* < 0.001).

**Figure 13.**
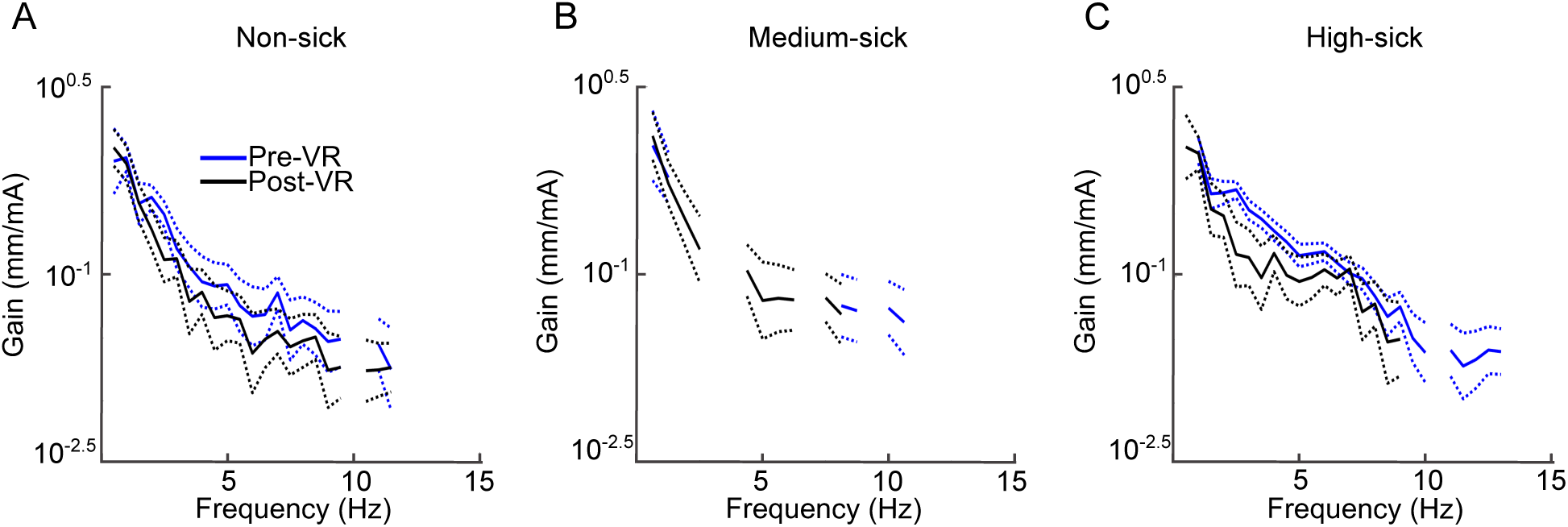
EVS-ML-CoP gain spectra during quiet-standing EVS trials for each sickness group. (A–C) Gain plotted as a function of frequency for the non-sick (A), medium-sick (B), and high-sick (C) groups, shown separately for pre-VR (blue) and post-VR (black) conditions. Gain is presented on a log scale. Solid lines represent group-averaged gain, and dotted lines indicate the 95% confidence limits, excluding frequencies without significant coherence for clarity.

### Gain

To complement coherence-based measures of vestibulomotor coupling, we examined EVS-ML-CoP gain to determine whether response amplitude followed similar patterns. No significant changes in EVS-ML-CoP gain from pre-to post-VR quiet-standing EVS trials were observed in the non-sick or medium-sick groups (Fig. 13A, B). Within the high-sick group, gain decreased significantly with a reduction of 64 ± 3% across 2.5–3.5 Hz (Fig. 13C).

Frequency regions in which the confidence limits did not overlap indicate significant differences between pre-and post-VR gain.

Between groups, EVS-ML-CoP gain was significantly greater in the high-sick group than in the non-sick group during the pre-VR quiet-standing EVS trials, with increases of 61 ± 2% across 3–4.5 Hz and 5.5–6.5 Hz (Fig. 14A) and remained elevated post-VR with a 72 ± 3% increase across 6–7 Hz (Fig. 14B).

**Figure 14.**
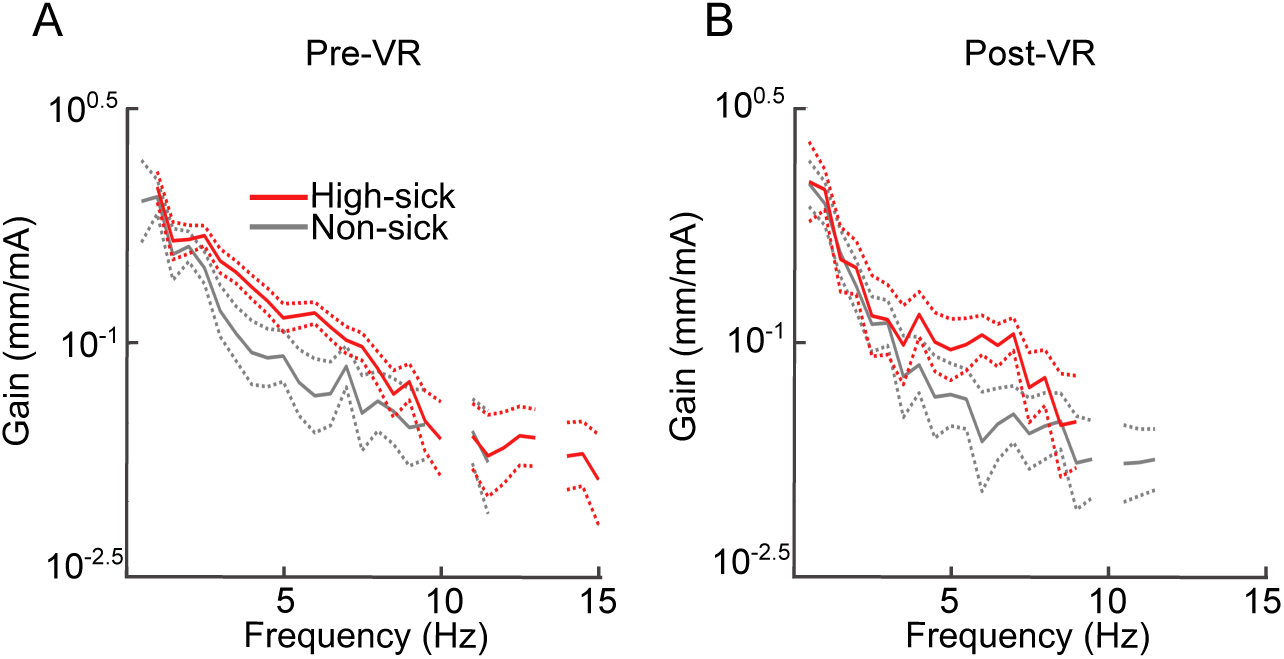
Comparison of EVS-ML-CoP gain spectra between non-sick and high-sick groups during quiet-standing EVS trials. (A, B) Gain plotted as a function of frequency for the non-sick (gray) and high-sick (red) groups during pre-VR (A) and post-VR (B) conditions. Gain is presented on a log scale. Solid lines represent group-averaged gain, and dotted lines indicate the 95% confidence limits, excluding frequencies without significant coherence for clarity.

When looking at first to second half changes during VR in EVS-ML-CoP gain, no significant differences were observed in the non-sick group (Fig. 15A). Within the medium-sick group, gain decreased significantly from the first to the second half of the VR trial, with a reduction of 57 ± 2% across 4–4.5 Hz (Fig. 15B).

**Figure 15.**
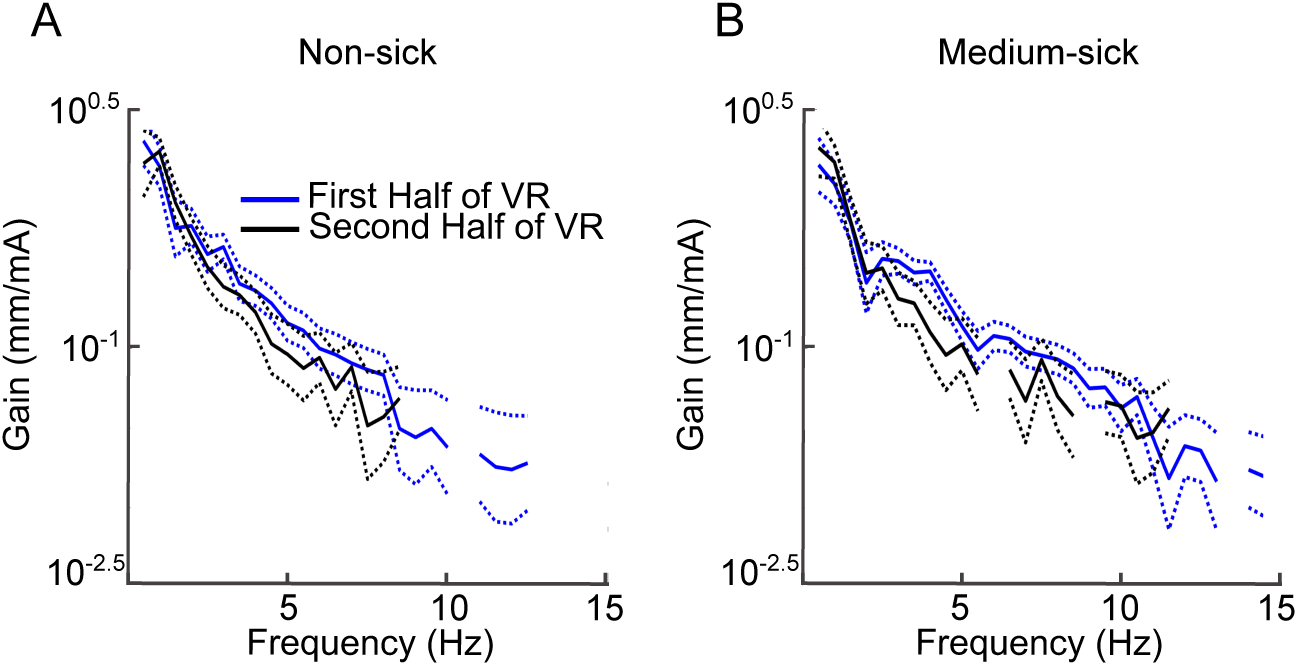
EVS-ML-CoP gain spectra during the VR trial for the non-sick and medium-sick groups. (A, B) Gain plotted as a function of frequency for the first half (blue) and second half (black) of the VR exposure in the non-sick (A) and medium-sick (B) groups. Gain is presented on a log scale. Solid lines represent group-averaged gain, and dotted lines indicate the 95% confidence limits, excluding frequencies without significant coherence for clarity.

### Post Hoc Analysis: New Sickness Groups

To assess the robustness of sickness-related differences in vestibulomotor coupling, coherence analyses were repeated using several alternative grouping strategies. Participants were first divided into tertiles based on FMS slope, with individuals exhibiting negative slopes (indicating symptom recovery) analyzed as a separate group. Tertiles were also generated after including participants with negative slopes. In addition, participants were grouped into tertiles based on total SSQ scores and, separately, based on pre-and post-VR EVS-ML-CoP coherence during quiet-standing, with corresponding FMS peak scores examined across these classifications. Across all alternative grouping strategies, the results were relatively consistent with those obtained using the primary peak-FMS classification (see Supplemental Material for detailed results). Coherence was the primary outcome of interest in these validation analyses, as it demonstrated the most robust and consistent effects. In contrast, ML-CoP RMS changes have been previously documented in the literature, EVS-ML-CoP gain analyses are inherently limited by the requirement for significant coherence.

### Post Hoc Analysis: FMS Slope Tertiles with Separate Negative Slope Group

In the FMS slope tertile classification with the separate negative slope group, participants were classified as tertile 1 (slope of FMS > 0 and ≥ 0.4; n = 12), tertile 2 (slope of FMS > 0.4 and ≤ 2; n = 11), tertile 3 (slope of FMS > 2; n = 11) and the negative slope group (slope of FMS negative; n = 4; Fig. 16).

**Figure 16.**
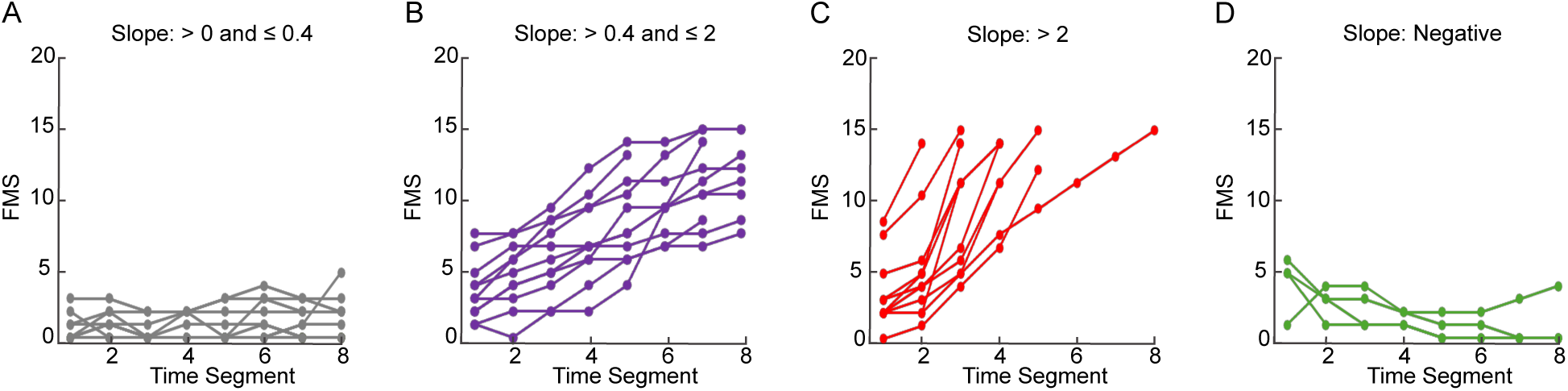
Participants grouped based on FMS slope values calculated across consecutive 2.5-minute time segments. (A–C) Participants grouped into tertiles based on positive FMS slope values. (D) Participants with negative FMS slopes. Each line represents an individual participant’s FMS trajectory across the VR trial.

Significant concatenated group-level EVS-ML-CoP coherence was observed across all groups (Fig. 17), with the largest frequency ranges in tertile 3 and the negative slope group (Fig. 17C, D).

**Figure 17.**
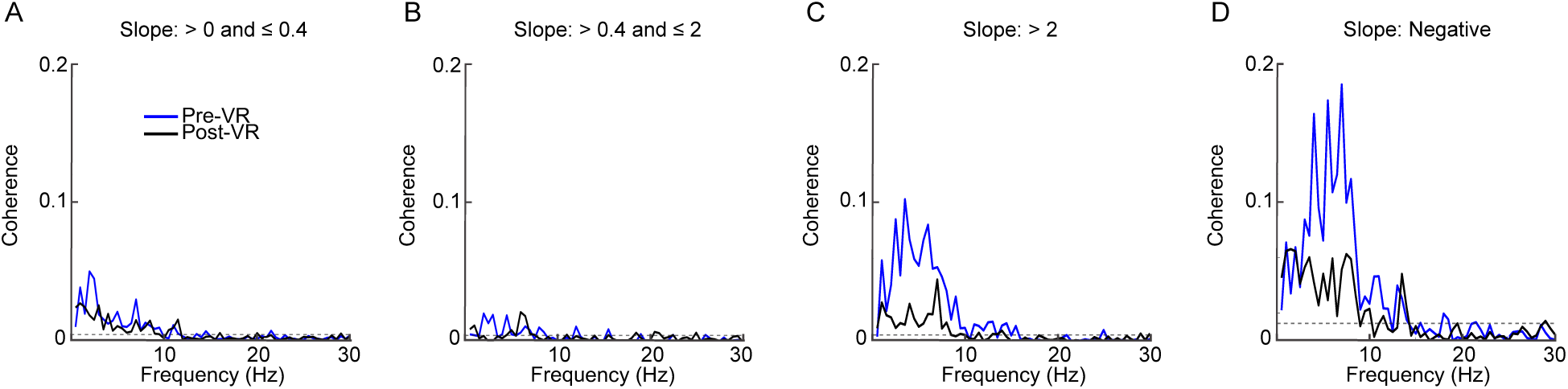
Concatenated group-level EVS-ML-CoP coherence spectra for participant groupings based on slopes of FMS trajectories. (A–D) Participants grouped based on individual FMS slope values calculated using linear regression across consecutive 2.5-minute VR time segments, showing tertile 1 (A), tertile 2 (B), tertile 3 (C), and the negative slope group (D). Coherence was plotted as a function of frequency for pre-VR (blue) and post-VR (black) quiet-standing EVS trials. A dashed horizontal line indicates the 95% confidence limit threshold for significant coherence.

Significant differences in concatenated group-level coherence between pre-and post-VR quiet-standing EVS trials were observed only in the third tertile (Fig. 18C). In this group, pre-VR coherence exceeded post-VR coherence across the 2.5–6.5 Hz frequency range. No significant pre-post differences in concatenated coherence were detected in the first or second tertiles, nor in the negative slope group (Fig. 18A, B, D).

**Figure 18.**
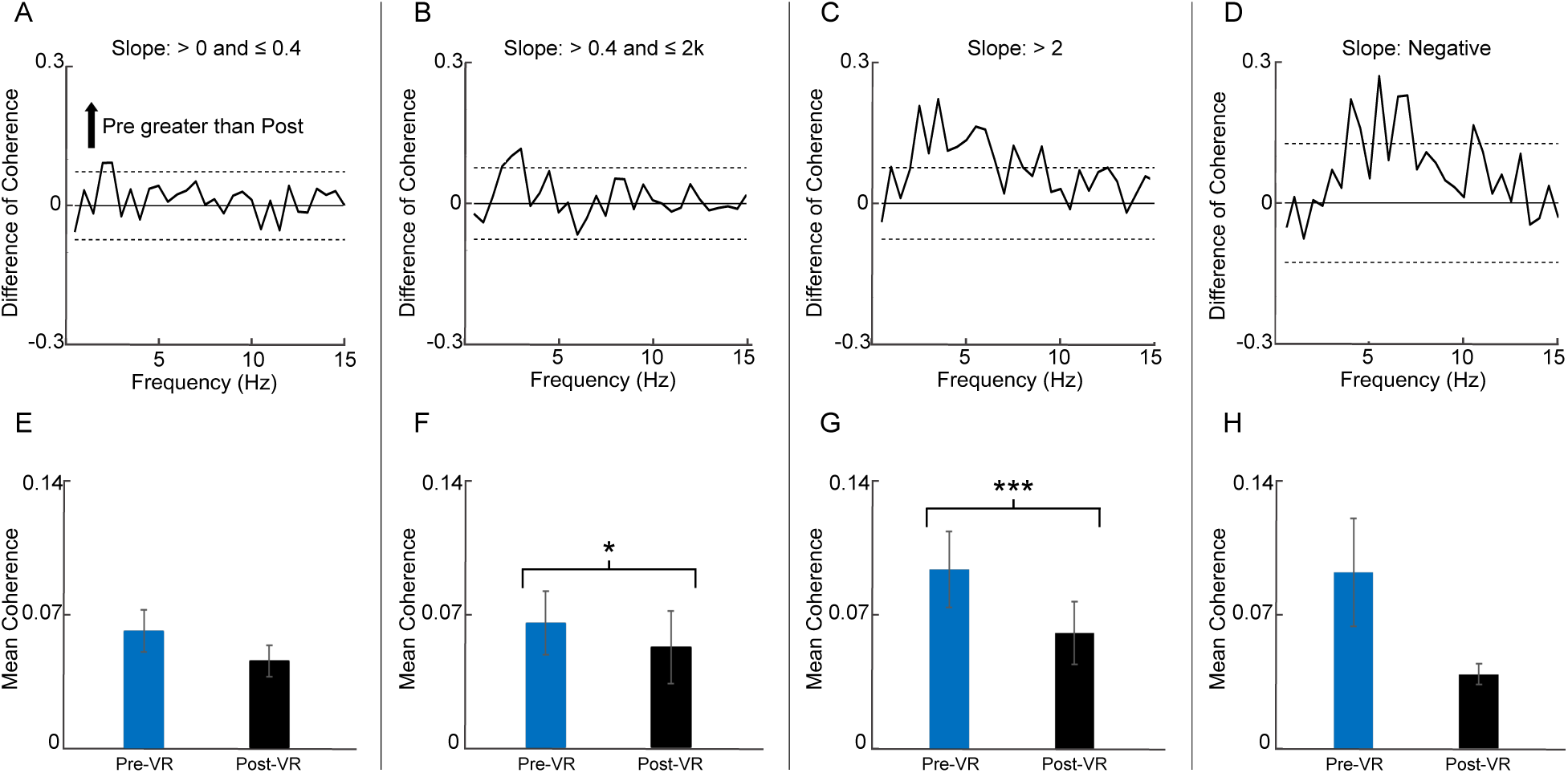
Pre-to post-VR changes in EVS-ML-CoP coherence across FMS slope-defined groups. (A–D) Frequency-specific difference-of-coherence tests for tertile 1 (A), tertile 2 (B), tertile 3 (C), and the negative slope group (D). Solid lines represent the difference in coherence across frequency, and horizontal lines indicate confidence limits. (E–H) Mean EVS-ML-CoP coherence averaged across 0–15 Hz and participants for pre-and post-VR quiet-standing EVS trials for tertile 1 (E), tertile 2 (F), tertile 3 (G), and the negative slope group (H). Bars represent group means and error bars indicate variability across participants. Asterisks indicate significant pairwise differences (* *p* < 0.05, ** *p* < 0.01, *** *p* < 0.001).

When coherence was averaged across frequencies from 0–15 Hz and across participants, the first tertile did not have significant changes pre-to post-VR (Fig. 18E). The second tertile showed elevated pre-VR coherence that decreased post-VR, reaching significance (Wilcoxon signed-rank test, *p* = 0.043, 𝑟_rb_ = 0.603; Fig. 18F). The third tertile also exhibited high pre-VR coherence, which declined significantly post-VR (repeated-measures *t*-test, *p* < 0.001, Cohen’s *d* = −1.50; Fig. 18G). The negative slope group was not included in statistical testing; descriptively, this group shows a decrease of −48.36 ± 11.32% (Fig. 18H).

Linear regressions assessing temporal changes in group-level coherence during the VR trial were significant for the negative slope group only, demonstrating the steepest decline, with a y-intercept of 0.0609 and a slope of −0.0043 (*p* = 0.013, R² = 0.67; Fig. 19C).

**Figure 19.**
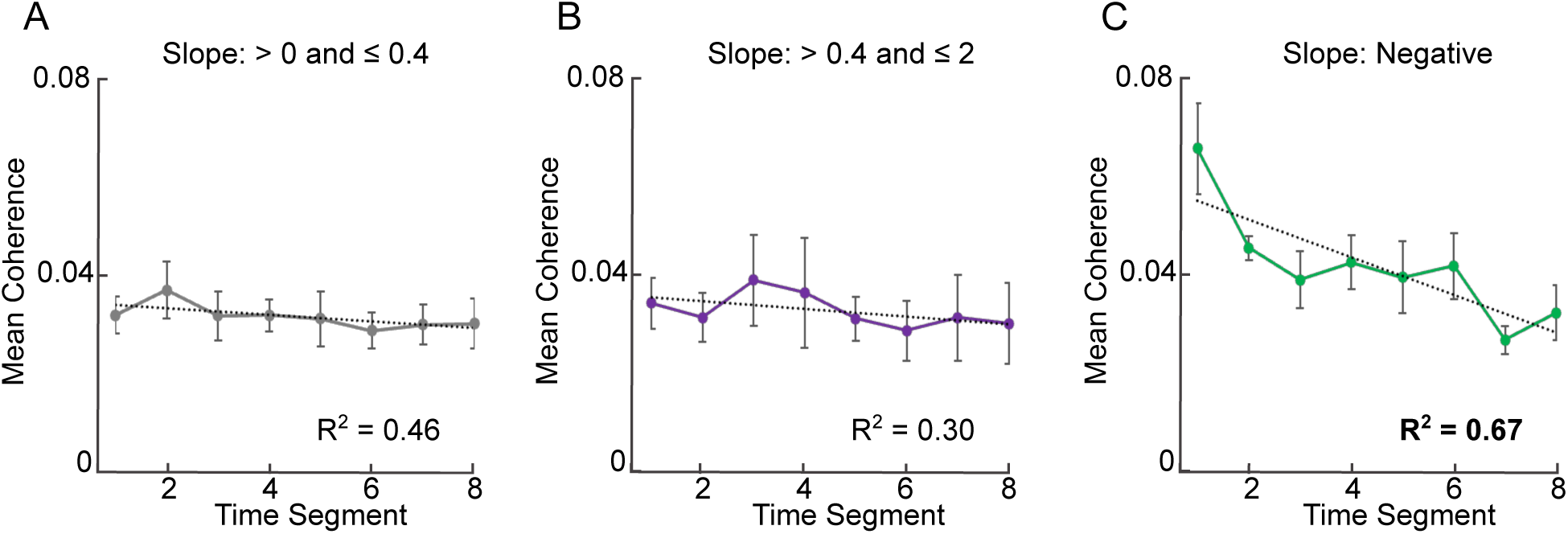
Group-averaged EVS-ML-CoP coherence across the VR exposure for tertile-based groupings derived from FMS slope classifications. (A–C) Mean EVS-ML-CoP coherence averaged across 0–15 Hz plotted across consecutive 2.5-minute time segments for tertile 1 (A), tertile 2 (B), and the negative-slope group (C). Points represent group means at each time segment, error bars indicate ± SEM, and dotted lines indicate best-fit linear regressions. R² values where *p* <0.05 are in bold.

## Discussion

This study examined how the CNS adjusts vestibulomotor weighting during immersive VR exposure with conflicting EVS and how these processes relate to individual susceptibility to cybersickness. We found that individuals who developed greater symptoms or terminated the VR exposure early exhibited elevated baseline vestibulomotor coupling (EVS-ML-CoP coherence) that declined over time, whereas non-sick individuals maintained consistently low coherence throughout exposure. Contrary to our initial hypothesis that greater vestibular reliance would mitigate sensory conflict, higher baseline vestibular weighting was associated with increased susceptibility to cybersickness. Across VR exposure, both coherence and gain decreased in symptomatic individuals, consistent with vestibulomotor downweighting. However, these reductions occurred alongside increases in postural sway, indicating that instability was not driven by increased vestibular contributions. Instead, this dissociation suggests that reweighting of vestibular input, although present, was insufficient or too slow to counteract conflicting sensory cues. Together, these findings highlight baseline sensory weighting as a key determinant of cybersickness susceptibility, with adaptive reweighting during exposure playing a secondary, mitigating role.

### Fast Motion Sickness and Simulator Sickness Scores Closely Related

Cybersickness severity differed across sickness groups in their SSQ measures (non-sick: 17.75 ± 3.35; medium-sick: 75.03 ± 15.1; high-sick: 105.66 ± 9.62). FMS and SSQ scores were strongly and linearly correlated, consistent with their validation as reliable measures of cybersickness (10,46). Among SSQ subscales, nausea showed the strongest association with peak FMS scores, exceeding associations with oculomotor and disorientation subscores, suggesting that nausea was the primary contributor to cybersickness severity in this study.

### Sickness Increases Postural Sway but Not Vestibular–Induced Sway

Our findings replicate previous work (23,27,56), demonstrating that postural sway (ML-CoP RMS) increases during VR exposure in individuals who develop greater cybersickness symptoms. ML-CoP RMS was greater in those with greater symptoms during post-VR quiet-standing EVS trials. In addition, symptomatic individuals exhibited larger increases in ML-CoP RMS from pre-to post-VR quiet-standing EVS trials (non-sick: −30.06 ± 7.67%; medium-sick:−7.00 ± 8.19%; high-sick: 29.18 ± 10.30%) and from the first to second half of the VR exposure.

Notably, the high-sick group exhibited substantially greater ML-CoP RMS during the first minute of the VR trial, which included the initial rollercoaster drop. This finding suggests that early postural responses to salient visual perturbations may serve as an indicator of susceptibility to severe cybersickness, with potential utility as a screening measure prior to extended VR exposure.

In contrast, vestibular-evoked sway showed an opposing pattern. EVS-ML-CoP gain was greater in the high-sick group at baseline (61 ± 6% higher than non-sick), indicating stronger vestibular-driven sway amplitude. Vestibular-evoked responses also decreased from pre-to post-VR quiet-standing EVS trials in the high-sick group (64 ± 6% reduction) and from the first to second half of the VR exposure in the medium-sick group (57 ± 2% reduction). This dissociation, where postural sway increased while vestibular-evoked responses decreased, indicates that increases in instability were likely not vestibular-driven. Instead, increases in instability may reflect increased reliance on visual information or other non-vestibular cues.

This pattern was not initially hypothesized, as vestibular cues were expected to be upweighted to compensate for unreliable visual motion (41). However, EVS introduces non-physiological and unreliable vestibular input, which is incongruent with natural head motion (29). Under these conditions, greater baseline reliance on vestibular signals may amplify sensory conflict rather than resolve it. Consequently, the observed downweighting of vestibular input over time likely reflects an adaptive response to reduce the influence of unreliable cues.

### Individuals Prone to Sickness Exhibit Greater Vestibulomotor Coupling, Which Decreases Over Time During VR

Consistent with gain findings, higher baseline coherence that declined over time was associated with greater symptoms. High-sick individuals exhibited the greatest initial coherence during pre-VR quiet-standing EVS trials, which declined post-VR (28 ± 7% decrease). Medium-sick participants also began with elevated coherence that decreased (35 ± 7% decrease). In contrast, non-sick participants maintained consistently low coherence across conditions, with a smaller decrease (19 ± 9% decrease). These patterns were preserved across alternative grouping strategies, indicating that the relationship between baseline coherence, its reduction, and sickness severity was robust to classification method.

At the group-spectral level, the medium-sick concatenated data appeared to exhibit particularly low coherence, reflecting high inter-individual variability rather than a true absence of vestibulomotor coupling. This interpretation is supported by the mean coherence results described above. Although concatenated coherence was greater in the high-sick group than in the non-sick group, no significant group differences were observed when coherence was averaged across frequencies (0–15 Hz) and participants. This discrepancy likely reflects substantial variability in coherence estimates across individuals and frequency bands.

Time-resolved analyses during VR exposure further support the interpretation that vestibulomotor coupling is initially elevated in symptomatic individuals and decreases over time. During the 20-minute VR trial, the medium-sick group exhibited higher initial coherence followed by a steady linear decline (mean slope = −0.0027) that was significantly steeper than that observed in the non-sick group. In contrast, the non-sick group showed minimal change over time (mean slope = −0.0005), maintaining consistently low coherence throughout exposure. The high-sick group also began with elevated coherence and exhibited a decline until trial truncation (slope = −0.006). These results are consistent with vestibular reweighting in response to sustained sensory conflict, although it remains unclear whether coherence would plateau with longer exposure.

Post hoc analyses indicated that a substantial portion of the negative slope observed in the medium-sick group was driven by a subset of participants who began with relatively higher FMS scores and exhibited symptom reductions over time. When analyzed separately, this group showed a steeper decline in coherence (mean slope = −0.0043), suggesting a greater capacity to downweight vestibular contributions during exposure, potentially contributing to symptom attenuation. This subgroup may reflect individuals who successfully adapt to sensory conflict through more rapid vestibular downweighting.

Together, these findings suggest that individuals who remain non-sick may not require substantial vestibular reweighting because their baseline vestibulomotor coupling is already low. In contrast, individuals who develop cybersickness begin with elevated vestibular reliance and downweight these signals over time, although this adaptation appears insufficient to prevent symptom progression.

### Limitations and Future Directions

Several limitations should be considered when interpreting the present findings. First, the study employed EVS with a peak amplitude of ±4.5 mA to probe vestibulomotor contributions to balance during VR exposure. While EVS provides a controlled method for quantifying vestibular influence, it introduces high-intensity, non-physiological and unreliable vestibular signals that likely increase cybersickness. Consequently, the observed reweighting reflects adaptation to perturbed vestibular input rather than to natural vestibular cues. Future studies could instead examine visuo-motor reweighting to better reflect more ecologically valid VR experiences.

Second, participant grouping was based on categorical classifications of sickness severity and task intolerance rather than continuous measures. While this approach captures meaningful functional differences (particularly the distinction between individuals who completed the task and those who terminated early) and facilitates frequency-dependent concatenation analyses, it may obscure more subtle relationships between symptom progression and sensory reweighting.

Third, the VR task was passive, meaning it did not require active head movement or user control, yet strongly challenged postural stability. As a result, the observed vestibulomotor responses reflect adaptation to strong externally imposed visual motion rather than self-generated movement. It remains unclear whether similar patterns of vestibulomotor weighting and reweighting would be observed in more interactive or self-directed VR environments, where predictability, agency, and sensorimotor contingencies differ.

Finally, the present findings suggest that baseline sensory reliance may be a key predictor of cybersickness susceptibility. Future work should examine whether baseline measures of vestibulomotor coherence or postural responses to brief visual perturbations can be used prospectively to identify individuals at risk for severe cybersickness. This approach could inform the development of personalized VR settings, adaptive training protocols, and targeted interventions to enhance tolerance and expand safe access to immersive technologies.

## Summary

In summary, baseline vestibulomotor weighting strongly influences susceptibility to cybersickness during VR exposure with EVS. Greater baseline coupling was associated with increased symptom severity and VR intolerance, whereas non-sick individuals maintained low vestibular contributions and required minimal reweighting. Although vestibulomotor reweighting occurred in symptomatic individuals, these changes were insufficient to prevent symptom escalation, highlighting baseline sensory reliance as a key determinant of vulnerability.

## Data Availability

All data and code for this article are provided in an Open Science Framework (OSF) repository, at https://osf.io/yw2t6. We report all measured variables and all analyses we conducted.

## Grants

This work was funded by Natural Sciences and Engineering Research Council of Canada (NSERC) Discovery Grant (RGPIN-03977-2020) to MBC.

## Disclosures and Disclaimers

The authors have no disclosures or disclaimers to declare.

## Author Contributions

MHG developed the main concept and theoretical framework of the project. MHG designed and conducted the experiment, analyzed the data, and led the writing of the manuscript. MBC secured funding and provided oversight of the project’s direction and planning. Both authors contributed critical feedback and helped shape the research, analysis, and manuscript.

