## Supplemental Materials for "Vestibulomotor Weighting Associated with Cybersickness in Virtual Reality"

**FMS Slope Tertiles**

In the FMS slope tertile classification with the separate negative slope group, participants were classified as tertile 1 (Slope of FMS > -0.6429 and ≤ 0.2; n = 13), tertile 2 (slope of FMS 0.2 and ≤ 1.9; n = 13) and tertile 3 (slope of FMS > 1,9; n = 12; Fig. S1).

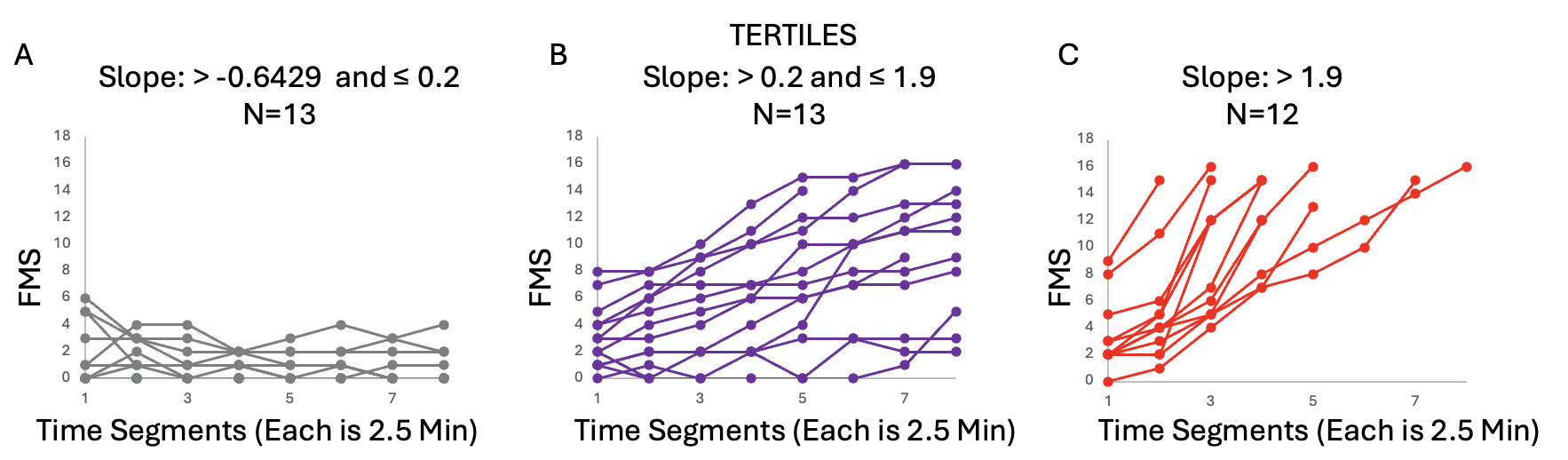
**Figure S1.** Participants grouped into tertiles based on individual FMS slope values calculated across consecutive 2.5-minute time segments. Panels show low (A), medium (B), and high (C) positive slope groups, with slope ranges and sample sizes indicated above each panel. Each line represents an individual participant’s FMS trajectory across the VR trial.

Significant concatenated group-level coherence was observed across all tertiles, with patterns broadly similar to those identified using the original peak FMS groupings. In the first tertile, significant coherence during pre-VR quiet-standing EVS trials was observed across 0.5–9.5 Hz, extending to 0.5–11.5 Hz post-VR. In the second tertile, significant pre-VR coherence was observed across 2–5 Hz and 6–7 Hz, with no significant coherence detected post-VR. In the third tertile, significant pre-VR coherence was observed across 1–10 Hz, 11–13 Hz, and 14–15.5 Hz, and post-VR coherence across 0.5–8.5 Hz and 13–14 Hz (Fig. S2).

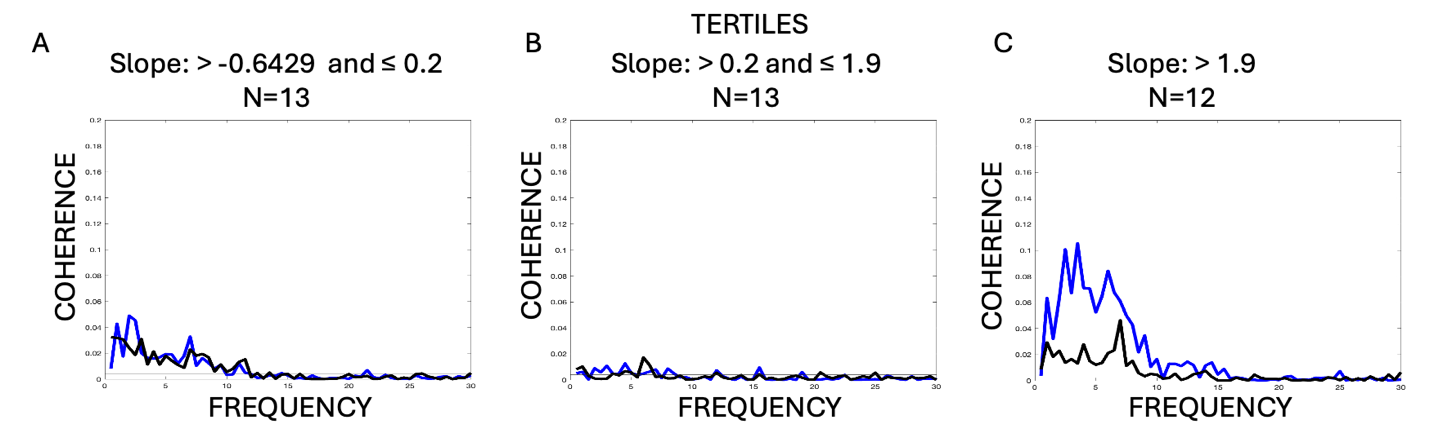

**Figure S2.** Concatenated group-level EVS–ML–CoP coherence spectra. Participants grouped into tertiles based on individual FMS slope values calculated using linear regression across consecutive 2.5-minute VR time segments, showing low (A), medium (B), and high (C) positive slope groups. Coherence is plotted as a function of frequency for pre-VR (blue) and post-VR (black) quiet-standing EVS trials. A dashed horizontal line indicates the 95% confidence limit threshold for significant coherence. Sample sizes and slope ranges are indicated above each panel.

Consistent with the original analysis, significant differences in concatenated group-level coherence between pre- and post-VR quiet-standing EVS trials were observed only in the third tertile (including participants with negative FMS slopes). In this group, pre-VR coherence exceeded post-VR coherence across the 2–6.5 Hz frequency range. No significant pre–post differences in concatenated coherence were detected in the first or second tertiles (Fig. S3).

When coherence was averaged across frequencies from 0–15 Hz and across participants, the third tertile exhibited the highest initial coherence during pre-VR quiet-standing EVS trials (0.104 ± 0.020), which declined significantly following VR exposure (post-VR: 0.066 ± 0.016; Wilcoxon signed-rank test, P = 0.021; rank-biserial correlation r = 0.645). The second tertile also showed elevated pre-VR coherence (0.063 ± 0.016) that decreased post-VR (0.052 ± 0.018), although this change did not reach significance. In contrast, the first tertile exhibited elevated pre-VR coherence (0.080 ± 0.013) that decreased significantly post-VR (0.050 ± 0.008; P = 0.015; Cohen’s d = −0.785). Despite these within-tertile changes, no significant differences in percent change of mean coherence from pre- to post-VR quiet-standing EVS trials were observed across tertiles (one-way ANOVA; tertile 1: −29.12 ± 7.53%; tertile 2: −22.76 ± 8.72%; tertile 3: −22.91 ± 14.73%; Fig. S3).

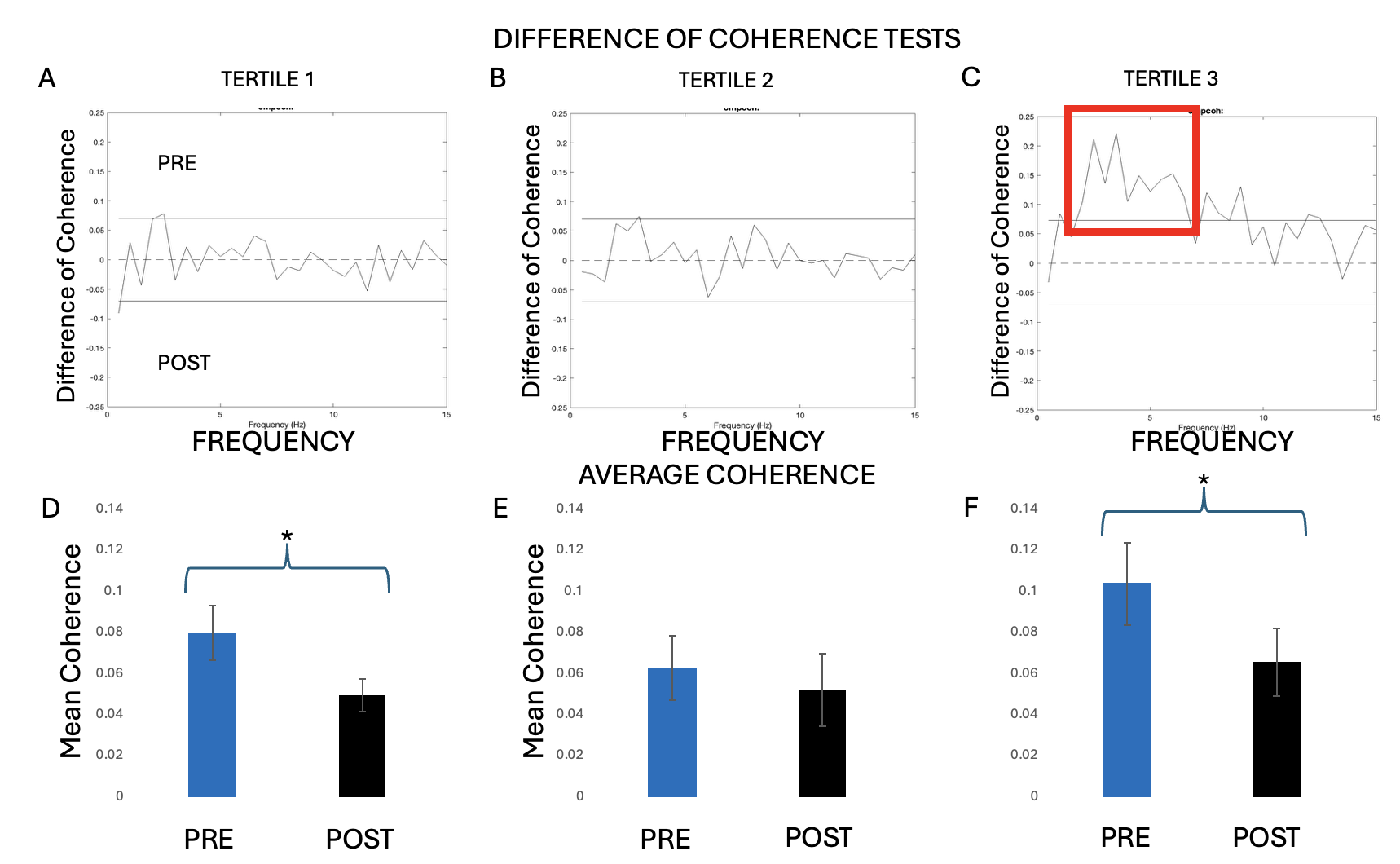

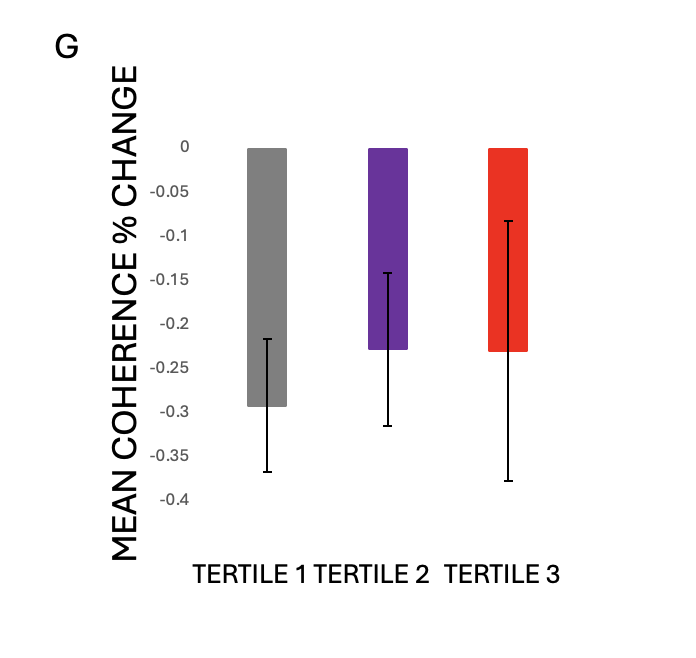

**Figure S3.** Difference-of-coherence and mean coherence measures during pre- and post-VR quiet-standing EVS trials across sickness groups. (A–C) Difference-of-coherence spectra comparing pre- and post-VR quiet-standing EVS trials for the non-sick (A), medium-sick (B), and high-sick (C) groups, plotted as a function of frequency. Horizontal lines indicate the 95% confidence limits used for the difference-of-coherence tests; frequency regions with nonoverlapping confidence limits are highlighted by red boxes. (D–F) Mean EVS–ML–CoP coherence averaged across frequencies (0–15 Hz) for pre-VR (blue) and post-VR (black) quiet-standing EVS trials in the non-sick (D), medium-sick (E), and high-sick (F) groups. Error bars represent ± SEM. (G) Percent change in mean EVS–ML–CoP coherence from pre- to post-VR quiet-standing EVS trials for each sickness group.

Significant differences in concatenated group-level coherence between the third tertile and first were observed during the pre-VR quiet-standing EVS trials, but not during the post-VR trials (like original). For these between-group coherence comparisons, an equal number of participants was required; therefore, one participant was removed from the first tertile group to match the third. In the pre-VR condition, coherence in the high-sick group exceeded that of the non-sick group across 2.5–8 Hz (same as original). When coherence was averaged across frequencies from 0–15 Hz and across participants to account for inter-individual variability (with no participants removed), Kruskal–Wallis tests revealed no significant differences across tertiles in either the pre- or post-VR quiet-standing EVS trials (Fig. S4).

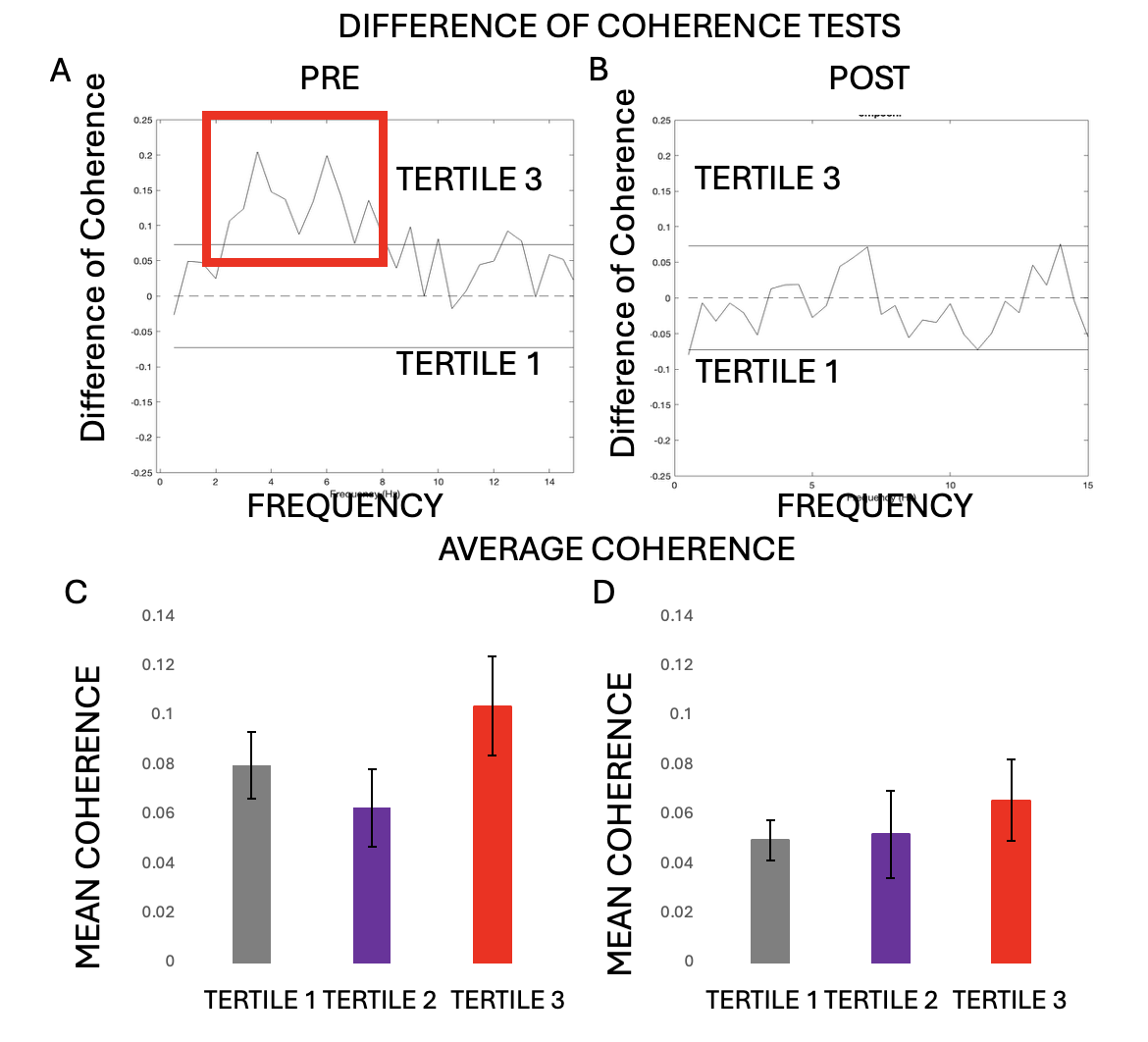

**Figure S4.** Coherence differences and averaged coherence across tertiles defined by FMS slope. (A–B) Difference-of-coherence tests comparing tertile 3 and tertile 1 during pre-VR (A) and post-VR (B) quiet-standing EVS trials, shown as the difference in coherence across frequency. Horizontal lines indicate confidence limits; red boxes highlight frequency ranges with significant differences. (C–D) Mean EVS–ML-CoP coherence averaged across frequencies from 0–15 Hz and across participants for each tertile during pre-VR (C) and post-VR (D) quiet-standing EVS trials. Bars represent group means and error bars indicate variability across participants.

Linear regressions assessing temporal changes in group-level coherence during the VR trial were significant for Tertiles 1 and 2. For Tertile 2, mean EVS–ML–CoP coherence averaged across participants for each 2.5-minute segment (eight segments total) yielded a y-intercept of 0.0373 and a negative slope of −0.0013 (R² = 0.65, P = 0.0151). Similarly, Tertile 1 exhibited a y-intercept of 0.0407 with a negative slope of −0.0015 (R² = 0.83, P = 0.00162; Fig. S5).

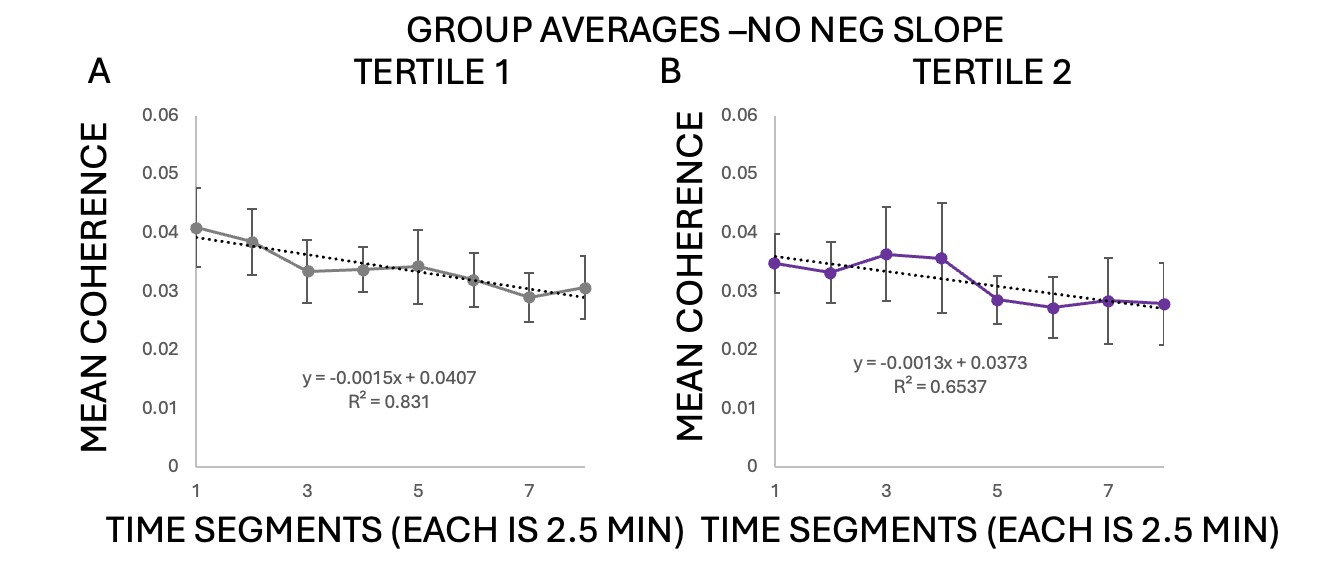
**Figure S5.** Group-averaged EVS–ML–CoP coherence across the VR exposure for tertile-based groupings derived from FMS slope classifications. Mean coherence (averaged across 0–15 Hz) is plotted as a function of time, with each time segment representing 2.5 min of VR exposure. (A–B) Group averages for Tertile 1 and Tertile 2 when participants with negative FMS slopes were not analyzed separately. Points represent group means at each time segment, error bars indicate ± SEM, and dotted lines denote best-fit linear regressions with corresponding equations and R² values displayed within each panel.

**SSQ Groups**

Tertile groups were defined based on total SSQ scores, and the corresponding FMS trajectories are shown in Fig. S6. The first SSQ tertile largely overlapped with the non-sick group, though it lost two participants to the medium-sick group and gained three from it. The second SSQ tertile closely resembled the medium-sick group but gained two participants from the non-sick group and five from the high-sick group, while losing three to the non-sick group and four to the high-sick group. The third SSQ tertile was most similar to the high-sick group, though it lost five participants to the medium-sick group and gained four from it.

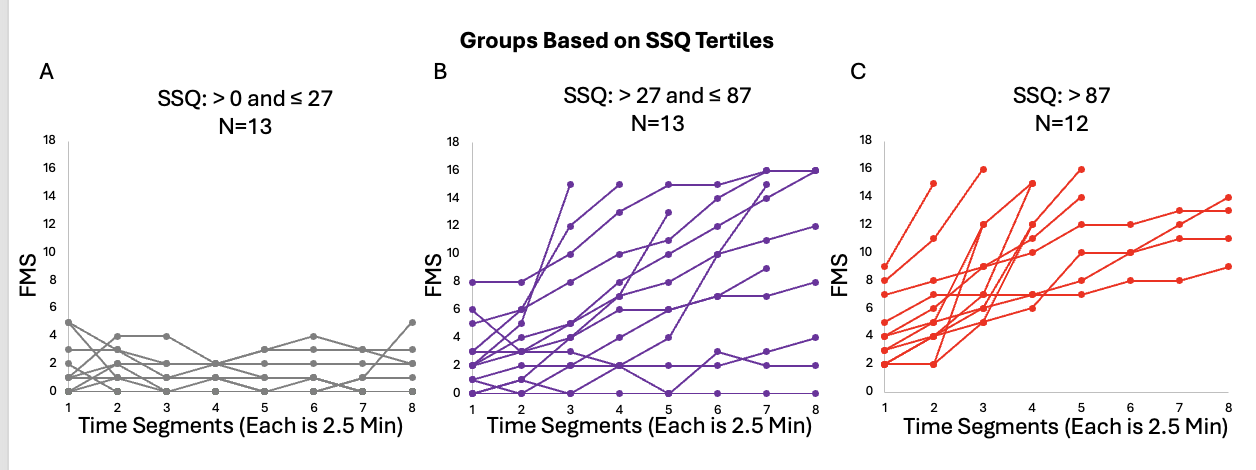

**Figure S6.** FMS trajectories during VR for groups defined by SSQ tertiles. Participants were grouped based on total SSQ scores into low-SSQ (A; SSQ > 0 and ≤ 27; N = 13), medium-SSQ (B; SSQ > 27 and ≤ 87; N = 13), and high-SSQ (C; SSQ > 87; N = 12) groups. Individual FMS ratings are shown across eight consecutive 2.5-minute time segments of VR exposure (total 20 minutes). Each line represents a single participant.

Significant concatenated group-level coherence was observed across all SSQ tertiles. In the first tertile, significant coherence during pre-VR quiet-standing EVS trials spanned 0.5–9.5 Hz. Post-VR coherence covered the same frequency range (0.5–9.5 Hz) and additionally extended to 10.5–11.5 Hz. In the second tertile, significant pre-VR coherence was observed across 0.5–10 Hz, 11–12.5 Hz, and 13.5–14.5 Hz, whereas post-VR coherence was present across 0.5–2 Hz and 3–8 Hz. In the third tertile, significant pre-VR coherence was observed across 2–8 Hz, with post-VR coherence spanning 0.5–2.5 Hz and 3.5–7 Hz (Fig. S7).

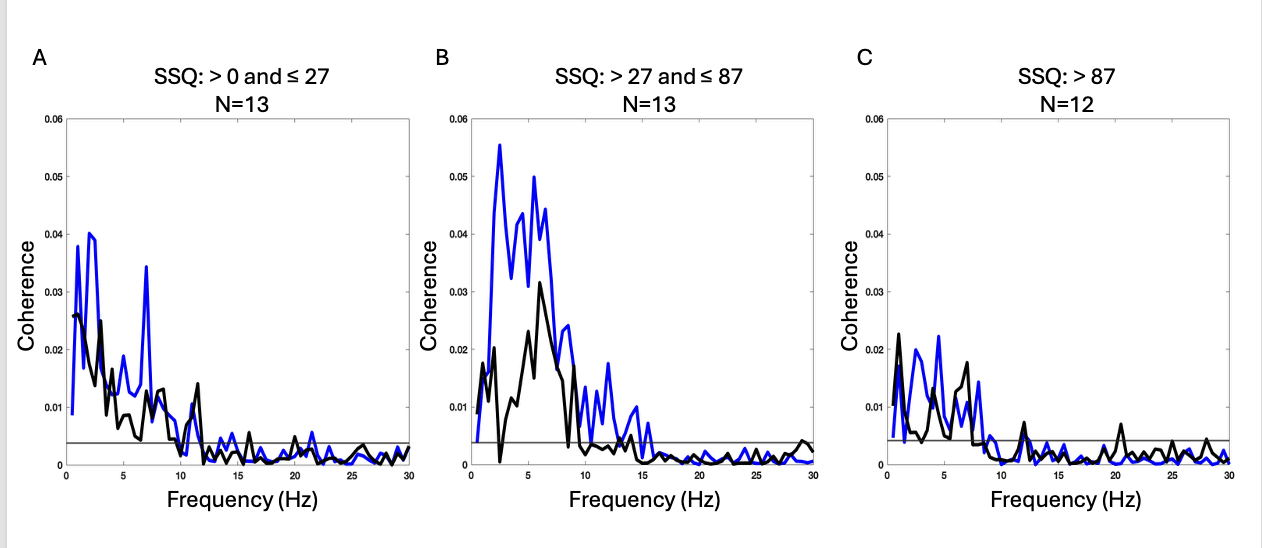

**Figure S7:** Concatenated group-level EVS–ML–CoP coherence spectra. Participants grouped into tertiles based on individual SSQ values, showing low SSQ (0 < SSQ ≤ 27; A), medium SSQ (27 < SSQ ≤ 87; B), and high SSQ (SSQ > 87; C) groups. Coherence is plotted as a function of frequency for pre-VR (blue) and post-VR (black) quiet-standing EVS trials. A dashed horizontal line indicates the 95% confidence limit threshold for significant coherence. Sample sizes and slope ranges are indicated above each panel.

Significant differences in concatenated group-level coherence between pre- and post-VR quiet-standing EVS trials were observed only in the second tertile. In this group, pre-VR coherence exceeded post-VR coherence across the 2.5–4.5 Hz frequency range. No significant pre–post differences in concatenated coherence were detected in the first or third tertiles.

When coherence was averaged across frequencies from 0–15 Hz and across participants, the second SSQ tertile exhibited the greatest initial coherence during pre-VR quiet-standing EVS trials (0.100 ± 0.021), which declined significantly following VR exposure (post-VR: 0.071 ± 0.020; Wilcoxon signed-rank test, *P* = 0.0061; rank-biserial correlation *r* = 0.717). The first tertile also showed elevated pre-VR coherence (0.071 ± 0.014) that decreased significantly post-VR (0.044 ± 0.007; *P* = 0.034; Cohen’s *d* = 0.664). Similarly, the third tertile exhibited elevated pre-VR coherence (0.073 ± 0.014) that declined significantly post-VR (0.052 ± 0.012; *P* = 0.0024; Cohen’s *d* = 0.804; Fig. S8).

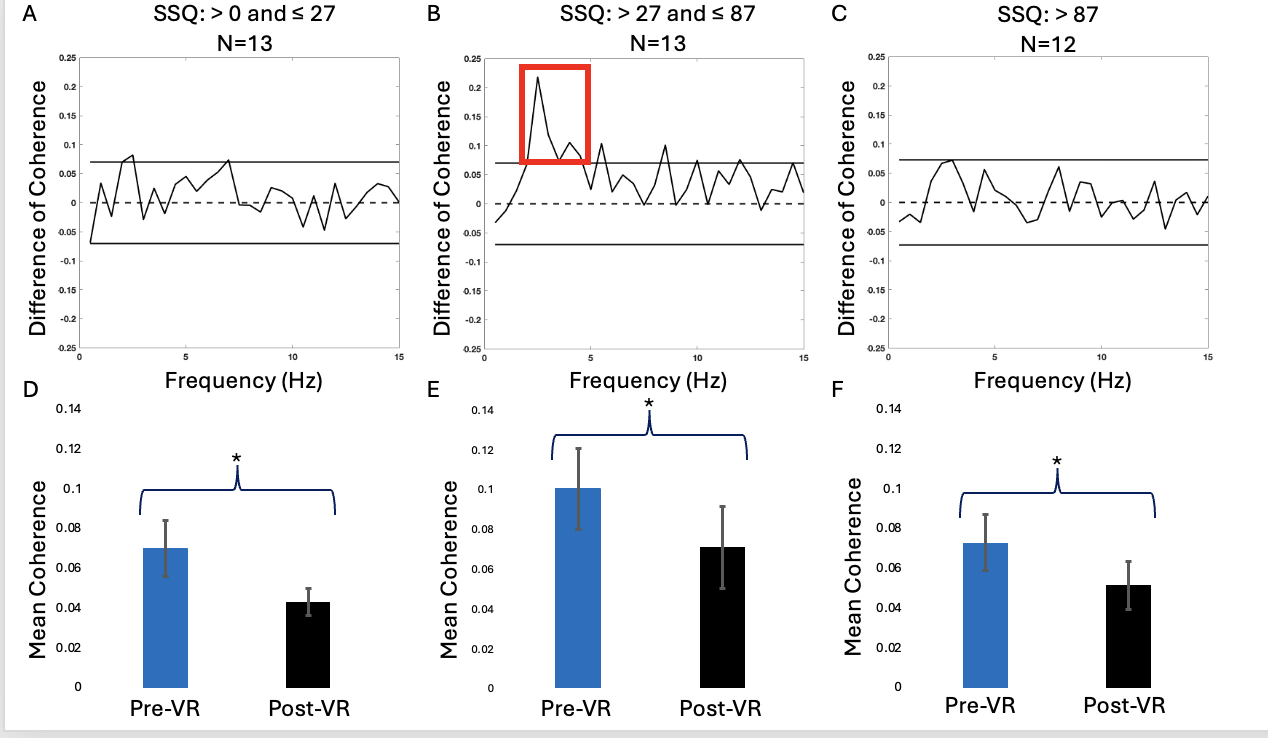

**Figure S8.** Changes in vestibulo-motor coherence across SSQ-based tertiles.
(A–C) Difference-of-coherence spectra comparing pre- and post-VR quiet-standing EVS trials for participants grouped by SSQ tertiles: low SSQ (0 < SSQ ≤ 27; A), medium SSQ (27 < SSQ ≤ 87; B), and high SSQ (SSQ > 87; C). Solid horizontal lines denote the 95% confidence limits, and the dashed line indicates zero difference. Red boxes highlight frequency ranges with significant differences. (D–F) Mean EVS–ML-CoP coherence averaged across 0–15 Hz and participants for pre- and post-VR quiet-standing EVS trials in the low (D), medium (E), and high (F) SSQ tertiles. Error bars represent ±1 SEM. Asterisks indicate significant pre- to post-VR differences within tertiles (p < 0.05).

SSQ-based grouping differed from FMS-based classification because SSQ scores reflect symptom severity on a comparatively coarse scale, with individual item ratings ranging from 1–3 rather than capturing continuous symptom progression over time. As a result, SSQ totals provide a less temporally sensitive measure of sickness severity than FMS, which tracks real-time symptom escalation during VR exposure. This difference in measurement resolution likely contributed to substantial overlap between SSQ tertiles and the original FMS-based sickness groups. In particular, considerable crossover was observed between participants originally classified as medium-sick and high-sick based on peak FMS scores. Consequently, SSQ-based groupings may be less precise in distinguishing gradations of sickness severity, especially for individuals whose symptoms evolved dynamically during exposure.

**Coherence Groups**

When participants were grouped into quartiles based on mean EVS–CoP coherence during the pre-VR quiet-standing trials, substantial overlap with the original sickness classifications was observed. Quartile 1 comprised 2 participants from the non-sick group, 6 from the medium-sick group, and 4 from the high-sick group. Quartile 2 included 8 non-sick, 3 medium-sick, and 2 high-sick participants. Quartile 3 consisted predominantly of high-sick participants, with 1 non-sick, 4 medium-sick, and 8 high-sick individuals (Table S1).

|  | Quartile 1 |  | Quartile 2 |  | Quartile 3 |
| --- | --- | --- | --- | --- | --- |
| Non-sick | 2 | Non-sick | 8 | Non-sick | 1 |
| Medium-sick | 6 | Medium-sick | 3 | Medium-sick | 4 |
| High-sick | 4 | High-sick | 2 | High-sick | 8 |

**Table S1.** Distribution of participants across coherence-based quartiles relative to original sickness group classifications. Participants were grouped into quartiles based on mean EVS–CoP coherence during pre-VR quiet-standing trials. Values indicate the number of participants from the original non-sick, medium-sick, and high-sick groups within each quartile, illustrating overlap between coherence-based grouping and sickness-based classification.

This table shows that baseline vestibulo-motor coherence does not map cleanly onto sickness-based groupings, but it is biased in a meaningful way. Quartile 3 (highest coherence) is enriched with high-sick participants (8 high-sick vs. 1 non-sick), suggesting that high baseline vestibulo-motor coupling is associated with greater sickness susceptibility, but is not sufficient on its own to predict sickness. Quartile 2 (mid coherence) contains the largest number of non-sick participants (8), implying that moderate coherence may be compatible with resilience, possibly because these individuals can downweight vestibular input effectively during VR. Quartile 1 (lowest coherence) is mixed, containing many medium-sick participants, indicating that low baseline coherence alone does not guarantee protection, and that other factors (e.g., visual reliance, reweighting capacity, or perceptual strategies) likely contribute. Overall, the overlap across quartiles supports the idea that baseline coherence is a risk factor rather than a categorical marker of sickness. It helps explain vulnerability trends at the group level but does not function as a one-to-one classifier.
